# Evolutionary patterns and repeated adaptive strategies of deep-sea anemones

**DOI:** 10.1101/2025.09.17.676462

**Authors:** Peidong Xin, Xiangxiang Wang, Yang Zhou, Chunhui Li, Wenjie Xu, Chenglong Zhu, Mingliang Hu, Yuxuan Liu, Ye Li, Jiangmin Zheng, Tao Qin, Yuan Yuan, Hui Shi, Yanjie Zhang, Liyan Qiao, Ping Li, Qiang Qiu, Kun Wang, Haibin Zhang, Chenguang Feng

**Author notes:** Contributed equally to this work (Peidong Xin, Xiangxiang Wang, Yang Zhou, Chunhui Li, Wenjie Xu, Chenglong Zhu).

## Abstract

Sea anemones occupy the full depth of the oceans, yet their evolutionary patterns and adaptive strategies to the enigmatic deep sea have remained contentious and poorly resolved. Here, we assemble genomes (*n* = 13) and transcriptomes for 15 species collected between 432 and 6,000 m and integrate them with all publicly available actiniarian data. Phylogenomic analyses reveal a mosaic topology among deep-sea and shallow-water clades. Using a novel framework that contrasts convergent gene-loss patterns, we show that a large number of light-associated gene families— including the complete circadian toolkit—were independently deleted after lineages entered the aphotic realm, whereas comparable loss in shallow taxa is negligible, providing decisive support for a shallow-water origin followed by multiple descents. Intriguingly, some deep-sea lineages further streamline energy budgets by recurrent loss or pseudogenisation of key meiotic genes (e.g., *Meiosin*, *Ythdc2*, *Spo11*, *Rad21*, *Mlh3*), indicating a shift towards asexual reproduction. Despite this extensive genomic erosion, deep-sea anemones exhibit sophisticated molecular tuning: specific amino-acid substitutions enhance protein stability and activity under deep-sea conditions, while selective expansions of gene families related to neural excitability, membrane systems, etc., likely mitigate the suppressive environmental effects on vital physiological processes. Enzyme activity assays in the yeast system confirm that the deep-sea variants exhibit superior activity and enhanced growth at 4°C. These results define a “loss-optimization-innovation” triad that underlies bathymetric adaptations and may apply to other deep-sea fauna worldwide.

## INTRODUCTION

Life in the deep sea, Earth’s largest yet least explored biome, endures perpetual darkness, crushing hydrostatic pressure, near-freezing temperatures, and severely limited food availability ^1–3^. Over the past decade, high-throughput sequencing and deep-sea sampling have yielded major genomic insights into deep-sea animals, including fishes and crustaceans ^4,5^. Yet, these studies focus on relatively derived lineages and leave the earliest stages of animal evolution under extreme conditions largely untested. As stemward representatives of Eumetazoan, sea anemones (Cnidaria: Anthozoa, Actiniaria) thrive from tide pools to the 10,000-m-deep hadal trenches ^6–9^; their simple, skeleton-less anatomy and well-documented plasticity make them uniquely tractable for disentangling cause-and-effect in adaptive change ^10^.

Consequently, investigating how actiniarians cope with deep-sea darkness, pressure, and cold is poised to bridge crucial gaps in our understanding of deep-sea adaptations in early-diverging multicellular animals, thereby offering clearer insights into the fundamental genomic principles that sustain life at our planet’s extremes.

Despite their ubiquity, two fundamental questions remain unresolved. First, where did sea anemones originate—shallow reefs or the abyss? The long-standing assumption of a shallow-water origin for most deep-sea fauna has rarely been put to a rigorous test. Isolated counter-examples, such as Campoy *et al.* ^11^ on cold-water corals, show that deep origins are possible. A morphology-plus-five-genes study by Rodríguez *et al.* ^12^ revealed striking convergence between deep- and shallow-dwelling actiniarians, yet the ancestral habitat of the order and the macro-evolutionary pattern of its bathymetric radiation remain unresolved, hampered by a sparse fossil record and extensive extinction ^12–15^. Second, regardless of directionality, which genomic mechanisms allow actiniarian lineages to traverse thousands of meters of depth?

Single-species genomes hint at confusing adaptive outcomes, including both reinforcement and breakdown of circadian systems, equivocal signals of positive selection, and idiosyncratic gene-family expansions ^16–19^, underscoring the absence of a coherent, order-wide framework.

To resolve these issues, we implemented a multi-pronged strategy integrating: (i) broad, depth-balanced genomic sampling, generating new genomes (*n* = 13) and transcriptomes (*n* = 15) from 15 species collected between 432 m and 6,000 m (Fig. 1; Supplementary Table 1), and merging these with all publicly available actiniarian datasets on genetics, taxonomy, distribution, and ecology; (ii) a phylogenomics-based test that infers ancestral habitat by contrasting convergent gene-loss patterns, a signal resilient to extinction bias; (iii) genome anatomy pinpoints genetic signatures associated with repeated adaptation, and (iv) functional assays that verify the physiological impact of deep-sea-specific variants. This comprehensive approach allowed us to revise actiniarian deep relationships, demonstrate a shallow-water origin followed by repeated descents, uncover systematic loss of meiotic and light-related genes that accompany a shift toward clonal reproduction, and validate protein-level adaptations that maintain respiration and membrane fluidity under high pressure and near-freezing temperatures, thereby revealing a unifying “loss-optimization-innovation” balance strategy of deep-sea adaptation.

**Fig. 1.**
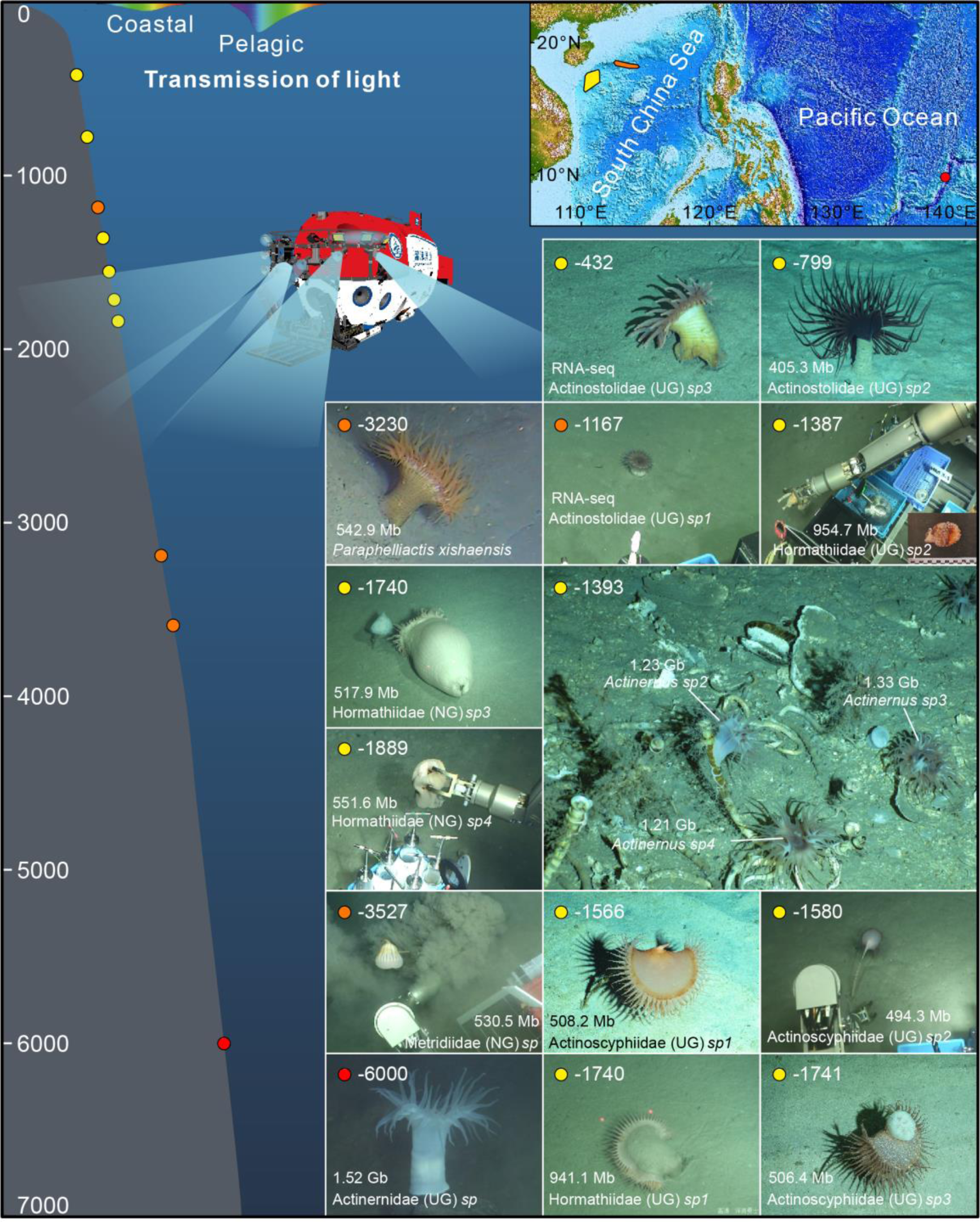
Sampling and sequencing information for deep-sea anemones. The color of the dots on the left side of the figure, representing different depths, corresponds to the color of the survey regions shown on the map in the upper right corner. The submersible depicted in the figure is the “*Shenhai Yongshi*” manned submersible, which served as the primary executor of the missions in this study.

## RESULTS AND DISCUSSION

### Repetitive DNA drives genome expansion in deep-sea anemones

This study provides a comprehensive genomic resource for 15 deep-sea anemone species (Fig. 1; Supplementary Tables 1-3). Using single-molecule real-time sequencing integrated with the BGI-seq platform, we generated high-quality genomes for seven representative species spanning the three major deep-sea superfamilies: Actinostoloidea, Metridioidea, and Actinernoidea (Supplementary Table 4). The resulting genome sizes ranged from 405.3 Mb to 1.52 Gb, consistent with *k*-mer-based estimates (Supplementary Table 4; Supplementary Fig. 1A). Assessment using BUSCO (metazoa_odb10) indicated high assembly completeness, ranging from 71.4% to 95.7% (Supplementary Table 4). These genomes were annotated with 27,110–32,245 predicted protein-coding genes, exhibiting gene set completeness (BUSCO protein recovery) ranging from 80% to 94.1% (Supplementary Table 5). Notably, these gene counts are comparable to those found in shallow-water anemone species (Supplementary Tables 5-6; Supplementary Fig. 1A). These quality metrics confirm that the generated genome assemblies and annotations are robust and suitable for downstream comparative and evolutionary analyses.

To further broaden our sampling, we performed deep whole-genome sequencing (161× to 542× coverage) for an additional six deep-sea species and generated transcriptomes for all 15 species to improve annotation and enable expression analysis (Supplementary Tables 2-4). We integrated these data with all publicly available actiniarian genomic (*n* = 13) and transcriptomic (*n* = 25) datasets (Supplementary Table 6), achieving comprehensive representation across all recognized superfamilies. This comparative analysis revealed a significant trend towards larger genome sizes in deep-sea species compared to their shallow-water congeners (Supplementary Fig. 1A), with genome size exhibiting a strong positive correlation with habitat depth within the deep-sea cohort (*p*-value = 3.6e-06; Supplementary Fig. 1B). This genome size expansion is primarily driven by the recurrent amplification of repetitive elements, rather than a proliferation of protein-coding genes (Supplementary Table 7; Supplementary Figs. 1A, C, D). This finding points towards the dynamic evolution of genome structure, particularly involving repetitive DNA, accompanying the diversification of anemones into the deep sea (Supplementary Fig. 1E).

### New superfamily phylogenetic relationship

Phylogenomic analysis of 1,849 high-quality single-copy orthologous genes from 41 species robustly recovered the monophyly of each of the five currently recognized superfamilies and supported the established subordinal classification dividing Actiniaria into Enthemonae and Anenthemonae (Fig. 2A; Supplementary Fig. 2). The topology derived from concatenated maximum likelihood analysis was largely congruent with that inferred using coalescent-based methods, with notable exceptions primarily among members of Actinernoidea (Fig. 2A; Supplementary Figs. 2, 3A).

**Fig. 2.**
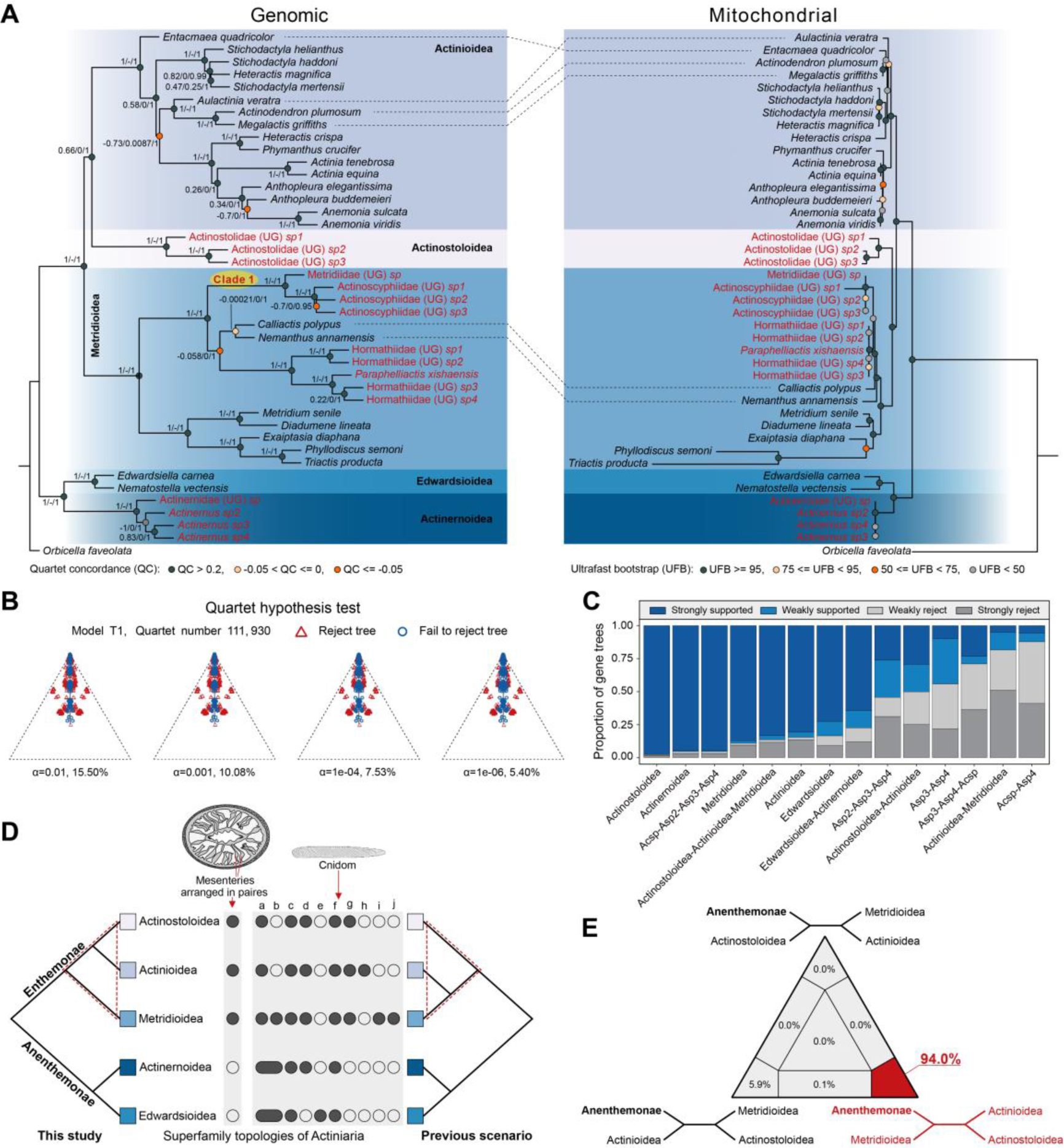
Phylogenomic analyses of deep-sea and shallow-water anemones. **(A)** The species tree (left) was inferred from 1,849 single-copy orthologous genes, while the mitochondrial gene tree (right) was reconstructed using 13 mitochondrial protein-coding genes. Taxa in red indicate deep-sea lineages. The values at each node represent the QC (Quartet Concordance) / QD (Quartet Differential) / QI (Quartet Informativeness) scores for internal branches of the species tree. QC > 0.2 represents strong support, while QC < 0 represents conflicting support. **(B)** Multispecies coalescent (MSC) analysis of the species tree inferred from ASTRAL. Each simplex plot represents the distribution of quartet concordance factors (qcCFs) under the T1 model (ILS) at a given significance level (α). Red triangles within the plot denote quartets that significantly reject ILS on the species tree. **(C)** Gene tree compatibility. Proportion of focal splits in gene trees that are strongly (or weakly) supported (or rejected) for the dispute genealogies. Abbreviations: Acsp: Actinernidae (UG) *sp*, Asp2: *Actinernus sp2*, Asp3: *Actinernus sp3*, Asp4: *Actinernus sp4*. **(D)** Phylogenetic framework of the five superfamilies within Actiniaria. The “previous scenario” refers to the superfamily relationships proposed by Rodríguez *et al.*^12^. Cnidom types a–j correspond to: a, spirocysts; b, basitrichs; c, holotrichs; d, microbasic p-mastigophores; e, atrichs; f, gracile spirocysts; g, microbasic b- and p-mastigophores; h, macrobasic p-mastigophores; i, robust and gracile spirocysts; j, p-amastigophores. **(E)** Likelihood mapping statistics showing the proportion of quartets supporting alternative placements of the three Enthemonae superfamilies. Quartets falling into the three triangle corners are considered informative. The red-labeled values denote the topology supported by this study and the corresponding proportion of support.

Both Quartet Concordance and Quartet Sampling analyses revealed significant gene tree incongruence and strongly conflicting signals across the phylogeny (Fig. 2A). Under the multispecies coalescent (MSC) model, 5.4% of quartets rejected the incomplete lineage sorting (ILS) model, even under a stringent significance threshold of α = 1e-6, suggesting that both ILS and interspecific gene flow have likely shaped the evolutionary history of sea anemones (Fig. 2B). Despite this underlying complexity, the coalescent-based topology represents the best-supported hypothesis of relationships (Fig. 2C; Supplementary Fig. 3) and is hereafter defined as the species tree.

While our species tree supports the traditional Enthemonae/Anenthemonae subdivision, it significantly challenges previous hypotheses regarding the relationships among the three superfamilies within Enthemonae (Fig. 2D). Contrary to the prevailing view placing Actinostoloidea as the sibling group to a clade uniting Actinioidea and Metridioidea ^12^, our analyses strongly support Metridioidea as the earliest diverging lineage within Enthemonae (Fig. 2D). Likelihood mapping analysis demonstrates that this topology is overwhelmingly supported by the underlying gene set (Fig. 2E), indicating that the conflict between previous perspective and our results likely stems from discordant signals between mitochondrial and nuclear datasets, rather than a lack of phylogenetic signal.

Inferring recent demographic histories using pairwise sequentially Markovian coalescent (PSMC) analysis revealed that nearly all examined sea anemones experienced substantial population declines during the Pliocene-Pleistocene transition (Supplementary Table 8; Supplementary Fig. 4A). Consistent with our previous observations ^19^, the onset of these declines generally occurred earlier in deep-sea species compared to their shallow-water counterparts (Supplementary Fig. 4B). This period is characterized by major marine extinctions, particularly affecting large-bodied fauna, such as mammals, seabirds, and sharks ^20,21^. It is particularly noteworthy that this environmental perturbation appears to have impacted anemone populations in the relatively stable deep-sea environment preferentially or at an earlier stage.

### Interrogation of ancestral origin status

The nested phylogenetic structure, with deep- and shallow-water lineages intermingled (Fig. 2A; Supplementary Fig. 2), strongly implies multiple independent transitions of sea anemones between these environments, indicative of repeated evolution. However, determining the predominant direction of these transitions – whether primarily shallow-to-deep or deep-to-shallow – requires resolving the ancestral habitat state of Actiniaria. Given the deep divergence times, significant lineage loss, and paucity of fossil evidence for sea anemones ^12–14^, traditional phylogenetic character state reconstruction methods are unlikely to yield robust conclusions. We therefore employed an alternative rationale: the probability of multiple independent lineages repeatedly losing the same gene is considerably higher than the probability of them repeatedly gaining the same gene *de novo*. By comparing the extent of convergent gene loss under alternative ancestral state scenarios, we can infer the most likely ancestral habitat and the primary direction of habitat transitions (Fig. 3A).

**Fig. 3.**
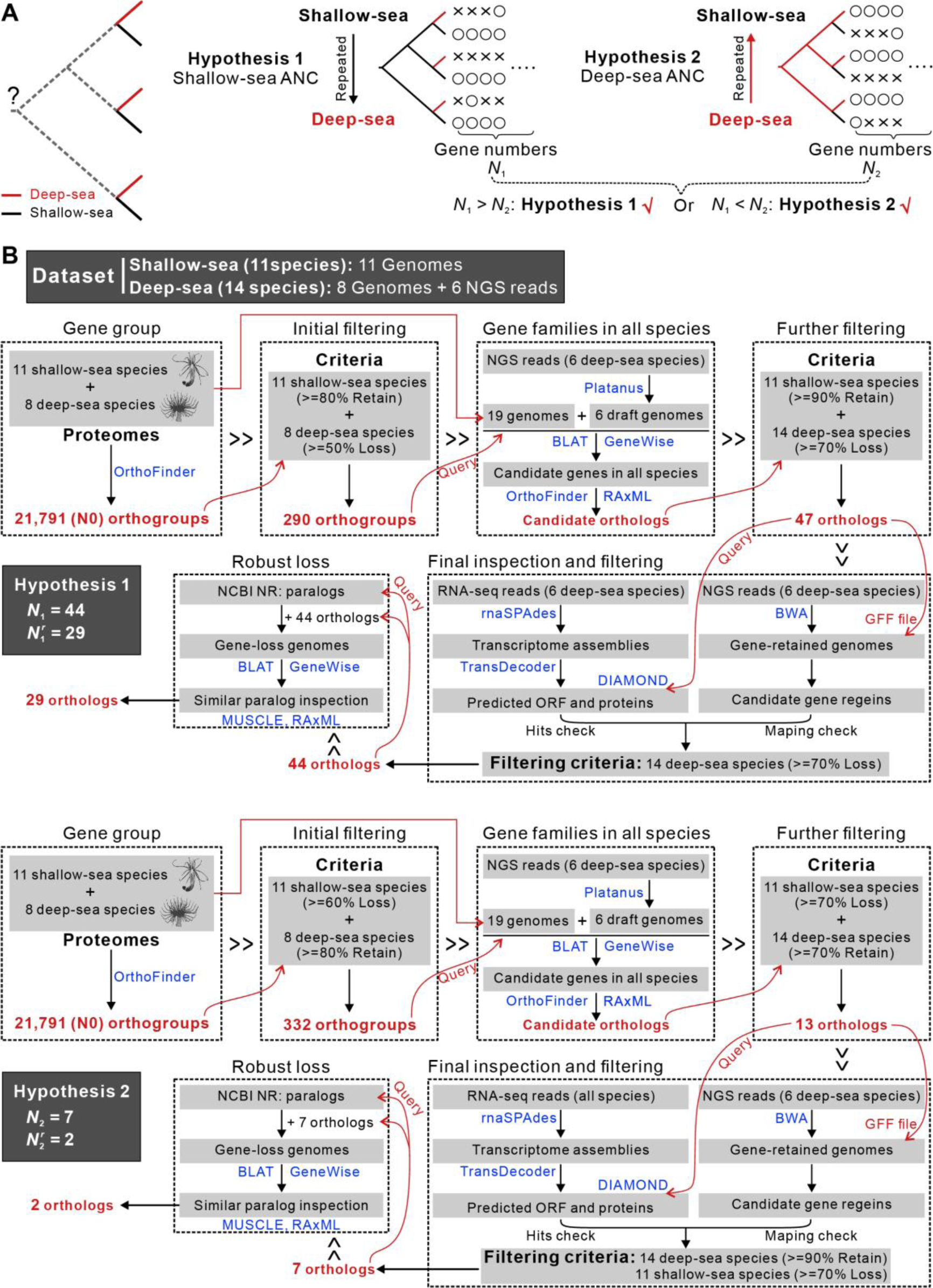
Hypothesis testing of evolutionary direction in sea anemones. **(A)** Schematic illustration of alternative hypotheses: the “Shallow-sea ANC” scenario assumes a shallow-water ancestral state, while the “Deep-sea ANC” scenario posits a deep-sea origin. The rationale of testing is that the probability of losing the same gene independently across multiple lineages is substantially higher than that of independent gene gain. Therefore, by assessing patterns of shared gene loss among lineages, we can infer the directionality of repeated habitat shifts and reconstruct the ancestral distribution of sea anemones. **(B)** Analytical framework and results under the two alternative hypotheses, based on patterns of recurrent gene loss.

Applying an optimized method for detecting repeated gene loss tailored to our dataset (Fig. 3B), we tested two competing hypotheses. Under the “Shallow-sea Ancestor (ANC)” hypothesis, we identified 44 gene families repeatedly lost across multiple deep-sea lineages; critically, 29 of these losses were deemed robust, lacking detectable paralogous backups in the respective genomes (Fig. 3B; Supplementary Table 9). Conversely, under the “Deep-sea ANC” hypothesis, only 7 gene families were repeatedly lost across shallow-water lineages, with merely 2 qualifying as robust losses (the remainder having paralogous backups) (Fig. 3B; Supplementary Table 10). This striking asymmetry provides compelling evidence favoring the “Shallow-sea ANC” hypothesis, suggesting that sea anemones originated in shallow waters and subsequently underwent multiple, independent colonizations of the deep sea.

Further examination of the genes repeatedly lost under the “Shallow-sea ANC” scenario revealed that they encompass nearly the entire molecular toolkit for the circadian clock pathway (Supplementary Table 9). The wholesale, repeated dismantling of such a deeply conserved and fundamental biological module ^22–26^ across disparate deep-sea lineages provides powerful, independent support for a shallow-water origin followed by repeated invasions of the deep, lightless environment.

### Repeated collapse of essential genes in deep-sea lineages

Our hypothesis testing revealed repeated losses of key functional genes across independent deep-sea lineages (Supplementary Table 9), which reflected a strategic streamlining, jettisoning functions unnecessary in the deep sea’s stable, resource-scarce environment. To gain a systematic understanding of the functional landscape shaped by gene loss, we performed a comprehensive survey across all species in our dataset, identifying not only repeatedly lost genes but also lineage-specific losses within relevant pathways or functional categories. This analysis targeted genes operating within the same pathways as those identified in the repeated loss screen, as well as genes with analogous functions. The results uncovered variable but significant patterns of gene loss within deep-sea anemones impacting crucial biological processes, including circadian rhythms, sensory perception, photoprotection, nutrition, and sexual reproduction.

#### Circadian rhythm machinery is largely dismantled

Before assessing circadian rhythm loss in deep-sea anemones, we first characterized the pathway architecture of the circadian rhythm in Actiniaria. Our analyses confirm the presence of at least seven core circadian components: *Cry1*, *Cry2*, *Cry-dash*, *Clock1*, *Clock2*, *Cycle*, and *Dec*. Notably, the *Per* gene appears restricted to Nephrozoa ^26–28^ (Supplementary Fig. 5). This composition suggests that the sea anemone circadian clock operates via two feedback loops regulated by *Cry* and *Dec* transcription factors, respectively (Fig. 4A) ^26,29^.

**Fig. 4.**
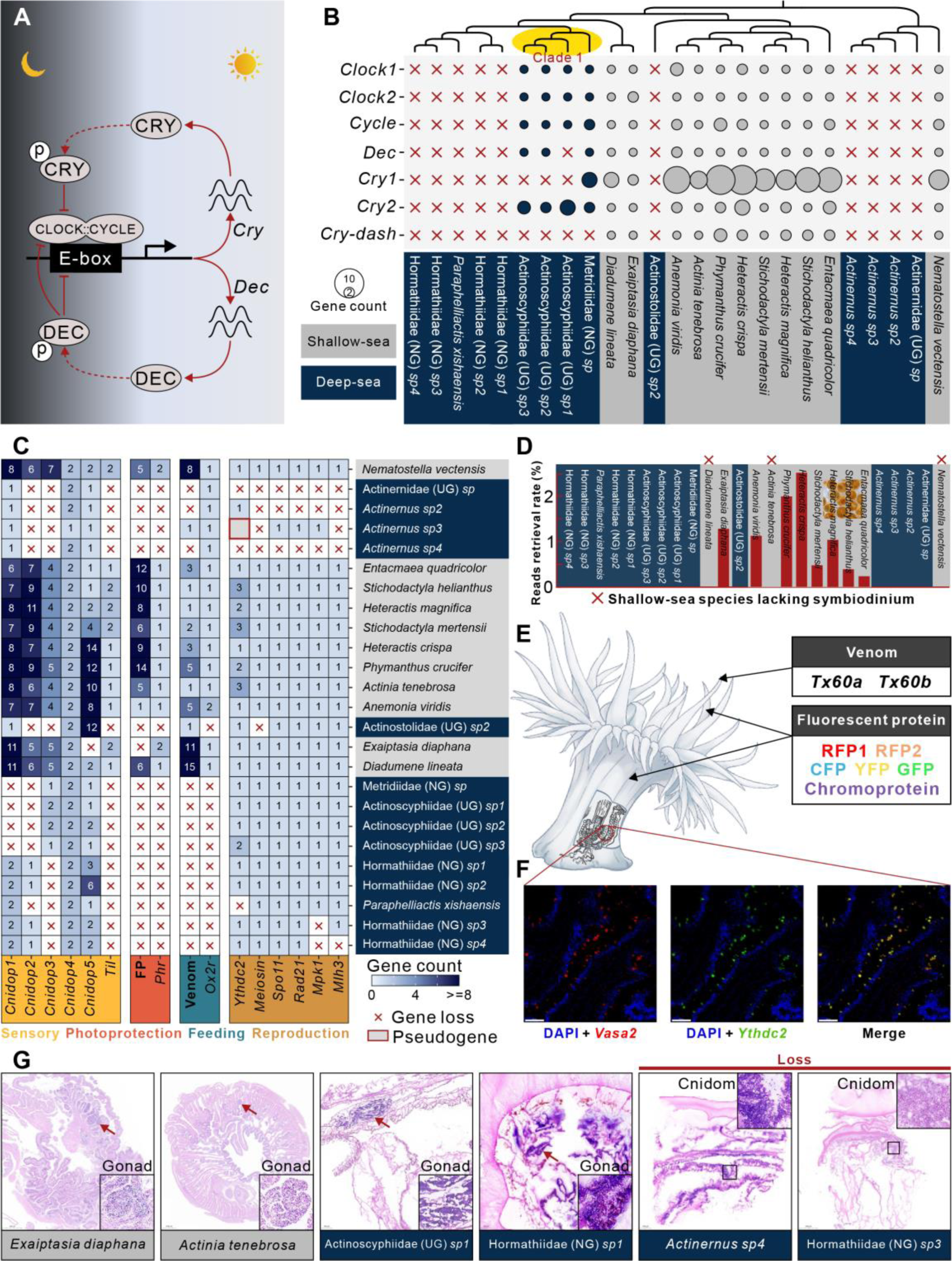
Repeated evolution of gene loss in deep-sea anemones. **(A)** A schematic diagram of the gene regulatory network for the circadian rhythm in sea anemones. **(B)** The number (circle size) and loss (red ‘X’) of circadian rhythm pathway genes in sea anemones. The yellow-highlighted Clade 1 corresponds to the red clade 1 in Fig. 2A. **(C)** The number and loss (red ‘X’) of genes related to functions of sensory, photoprotection, feeding, and reproduction in sea anemones. The “FP” refers to fluorescent proteins. The “Venom” refers to two toxin genes, *Tx60a* and *Tx60b*. **(D)** Assessment of symbiotic algae in sea anemones. The x-axis represents species, and the y-axis shows the effective retrieval rate of NGS reads against to ITS2 database. “X” indicates species known to lack endosymbiosis. A blue background represents deep-sea lineages, while a gray background represents shallow-water lineages. **(E)** Diagram of the expression locations of the venom genes and Fluorescent proteins in sea anemones. **(F)** *In situ* hybridization results for the *Vasa2* and *Ythdc2* genes. The sampled region is the columnar tissue of the gonadal area in *Exaiptasia diaphana* (refer to Fig. 4E). **(G)** Histological HE staining of the gonadal area in deep-sea and shallow-water anemones. Enlarged views of arrow-marked regions are shown in insets. The background color of the species names indicates their distribution: gray for shallow-sea species and blue for deep-sea species. The results show that no gonadal tissue was observed in *Actinernus sp4* and Hormathiidae (NG) *sp3*.

Genome-wide surveys revealed that shallow-water lineages consistently retain the full complement of these circadian genes (Fig. 4B). In stark contrast, nearly all deep-sea species examined, with the specific exception of those within Clade 1 of the superfamily Metridioidea, have lost the entire circadian pathway toolkit (Fig. 4B).

This wholesale loss reflects the absence of predictable light-dark and temperature cycles in most deep-sea habitats. Intriguingly, although species in Metridioidea Clade 1 exhibit some reduction in circadian gene copy number compared to shallow relatives, they retain both the *Cry*- and *Dec*-based feedback loops, and all constituent genes remain actively transcribed (Fig. 4B; Supplementary Fig. 6), indicating functional rhythm preservation. The four Clade 1 species in our study inhabit depths between 1,566–3,527 m (Fig. 1), well within the aphotic, thermally stable zone ^30^.

Therefore, the retention of circadian machinery in Clade 1 appears to be a lineage-specific trait, independent of ambient light or temperature cues at their collection depths. Given that all known members of this clade exhibit pelagic dispersal phases or lifestyles influenced by ocean currents^31–34^, their retained rhythmicity might be linked to this unique ecological trait.

#### Sensory modifications for a lightless, stable world

We next investigated sensory system modifications in deep-sea anemones. Repeated loss analyses had already implicated genes involved in thermosensation (*Til*) and photosensation (*Cnidop2*, *Cnidop3*). Our broader genomic survey confirms the specific loss of *Til*, a key gene for detecting temperature fluctuations ^35^, across all examined deep-sea lineages (Fig. 4C). The opsin gene family, responsible for light detection, comprises five members in sea anemones: *Cnidop1*, *Cnidop2*, *Cnidop3*, *Cnidop4*, and *Cnidop5*. Among these, *Cnidop1*, *Cnidop2*, and *Cnidop3* show marked contractions or complete loss in multiple deep-sea lineages (Fig. 4C). Consistent with compromised photosensitivity, *in situ* observations during sample collection failed to detect any behavioral response of deep-sea anemones to the lights of our remotely operated vehicle (Supplementary Videos 1-6). We also specifically examined the repertoire of Transient Receptor Potential (TRP) channels, a conserved metazoan channel family involved in diverse sensory modalities including vision, olfaction, mechanosensation, and nociception ^36–38^. Anemones possess seven TRP subfamilies (*Trpa*, *Trpc*, *Trpm*, *Trpml*, *Trpv*, *Trpvl*, and *Trpp*), but unlike opsins, the gene content within these subfamilies appears remarkably stable across both shallow and deep lineages, with no evidence for major lineage-specific expansions or contractions (Supplementary Fig. 7).

Overall, adaptation of sea anemones to the deep sea involved specific, albeit somewhat limited, modifications to the sensory toolkit, primarily entailing the reduction or complete loss of light and temperature fluctuation sensing capabilities.

#### Loss of photoprotection and shifts in nutrition

In the perpetual darkness of the deep sea, photoprotective mechanisms essential for surface life become redundant. Accordingly, we observed the complete loss of the *Phr* (Photolyases) gene, crucial for repairing UV-induced DNA damage ^39,40^, in all examined deep-sea anemones (Fig. 4C). Concurrently, genes encoding fluorescent proteins (FPs), which can serve photoprotective roles in cnidarians ^41–43^, are conspicuously absent from all deep-sea lineages (Fig. 4C). While the lack of vibrant coloration in deep-sea anemones is partly due to the absence of photosynthetic symbionts (see below), the loss of FPs represents a direct genomic contribution to this phenotype.

The phototactic behavior common in shallow-water anemones is often linked to the photosynthetic needs of their algal symbionts ^41,44–46^. Given the absence of light, do deep-sea anemones lack algal symbionts? To address this question, we screened our whole-genome sequencing reads against a comprehensive database of symbiont Internal Transcribed Spacer 2 (ITS2) tags ^47–49^. This yielded zero hits for ITS2 sequences in all deep-sea species (Fig. 4D), strongly suggesting that endosymbiosis with photosynthetic algae is indeed absent in these lineages.

Symbiotic algae in shallow-water anemones are well known to provide essential nutrients to their hosts, promoting growth and reproduction ^50–54^; their absence implies fundamental shifts in the nutritional ecology of deep-sea anemones. Supporting this, we found that genes encoding two toxins, *Tx60a* and *Tx60b*, both localized to the predatory tentacles ^55^, are lost in the vast majority of deep-sea lineages while being retained in all shallow-water relatives (Fig. 4C, E; Supplementary Fig. 8). As sea anemone toxin repertoires are known to correlate strongly with prey composition ^56–58^, the loss of these specific toxins likely reflects alterations in diet or feeding strategy in the deep sea. Furthermore, the gene *Ox2r*, implicated in appetite and feeding regulation ^59,60^, is also convergently lost across deep-sea lineages (Fig. 4C). Collectively, these findings point towards profound shifts in the feeding and nutritional strategies of anemones upon colonizing the deep sea.

#### Erosion of sexual reproduction pathways

For sea anemones capable of both sexual and asexual reproduction, nutritional and environmental conditions play a crucial role in determining which reproductive strategy is adopted ^61,62^. The findings above revealed a series of responses by sea anemones to environmental and nutritional changes following their colonization of the deep sea. To further investigate whether these changes have influenced the reproductive strategies of anemones, we first examined the gene *Vasa*, which is specifically expressed in germline progenitor cells and serves as a marker gene for pedal laceration during asexual reproduction in anemones ^63,64^ (Fig. 4F). The results showed that all investigated anemones retained both *Vasa1* and *Vasa2* (Supplementary Fig. 9).

Next, we analyzed the meiotic toolkit and revealed significant gene degradation in deep-sea lineages within the superfamilies Actinostoloidea, Metridioidea, and Actinernoidea (Fig. 4C; Supplementary Table 11). Key genes essential for distinct meiotic stages show evidence of loss or pseudogenization, including those critical for meiotic initiation (*Meiosin*; *Ythdc2*, Fig. 4F), homologous chromosome pairing (*Spo11*), sister chromatid cohesion (*Rad21*), Meiosis I progression (*Mpk1*), and Meiosis II completion (*Mlh3*). These losses cripple gamete production, curtailing sexual reproduction ^65–71^. Loss events were particularly pronounced within the Actinernoidea lineage (Fig. 4C). Histological evidence confirms this shift: species with degraded meiotic genes lack gonads, their tissue regions occupied by cnidom (nematocyst clusters), unlike those with intact meiotic genes (Fig. 4G). Thus, some deep-sea anemones have streamlined their biology, favoring energy-conserving asexual propagation.

### Protein optimization and expansion facilitate adaptability

Adaptation to the deep sea is not solely a subtractive process of gene loss; it also involves constructive changes, namely the molecular optimization of existing proteins and the strategic expansion of specific gene families. We investigated both phenomena to obtain a complete picture of adaptive strategies.

#### Optimized proteins counteract deep-sea challenges

The high hydrostatic pressure and low temperatures characteristic of the deep sea pose severe challenges to protein stability and enzymatic activity ^4,72,73^. Consequently, adaptive modifications at the amino acid level are expected. However, the vast evolutionary timescale covered by our study (∼434 Ma; Supplementary Fig. 10) results in near-saturation of substitutions within many gene lineages (Supplementary Fig. 11), precluding the reliable application of standard selection pressure analyses (e.g., dN/dS) to detect positive selection or rapid evolution. To circumvent this limitation, we implemented a simplified analytical pipeline focused on identifying deep-sea lineage-specific amino acid variants within functionally conserved protein regions, leveraging information entropy metrics (Supplementary Fig. 12).

A total of 169 genes were identified with lineage-specific amino acid substitutions present in all deep-sea lineages (Supplementary Table 12), termed Deep-sea Specific Genes (DSGs). DSGs are defined by amino acid substitutions that are conserved within each deep-sea lineage but not necessarily conserved across different lineages. These DSGs are involved in fundamental cellular processes, including metabolism (e.g., *Ndufa12*), transcription (e.g., *Elob*), and protein folding (e.g., *Hsp90b*) (Fig. 5A-B; Supplementary Fig. 13), implicating systemic physiological adjustments to the deep-sea environment. Metabolic functions were particularly enriched, accounting for 23.6% of identified DSGs (Fig. 5A). Notably, all five complexes of the mitochondrial respiratory chain contained constituent genes identified as DSGs (Fig. 5C), underscoring the importance of energy metabolism adaptation.

**Fig. 5.**
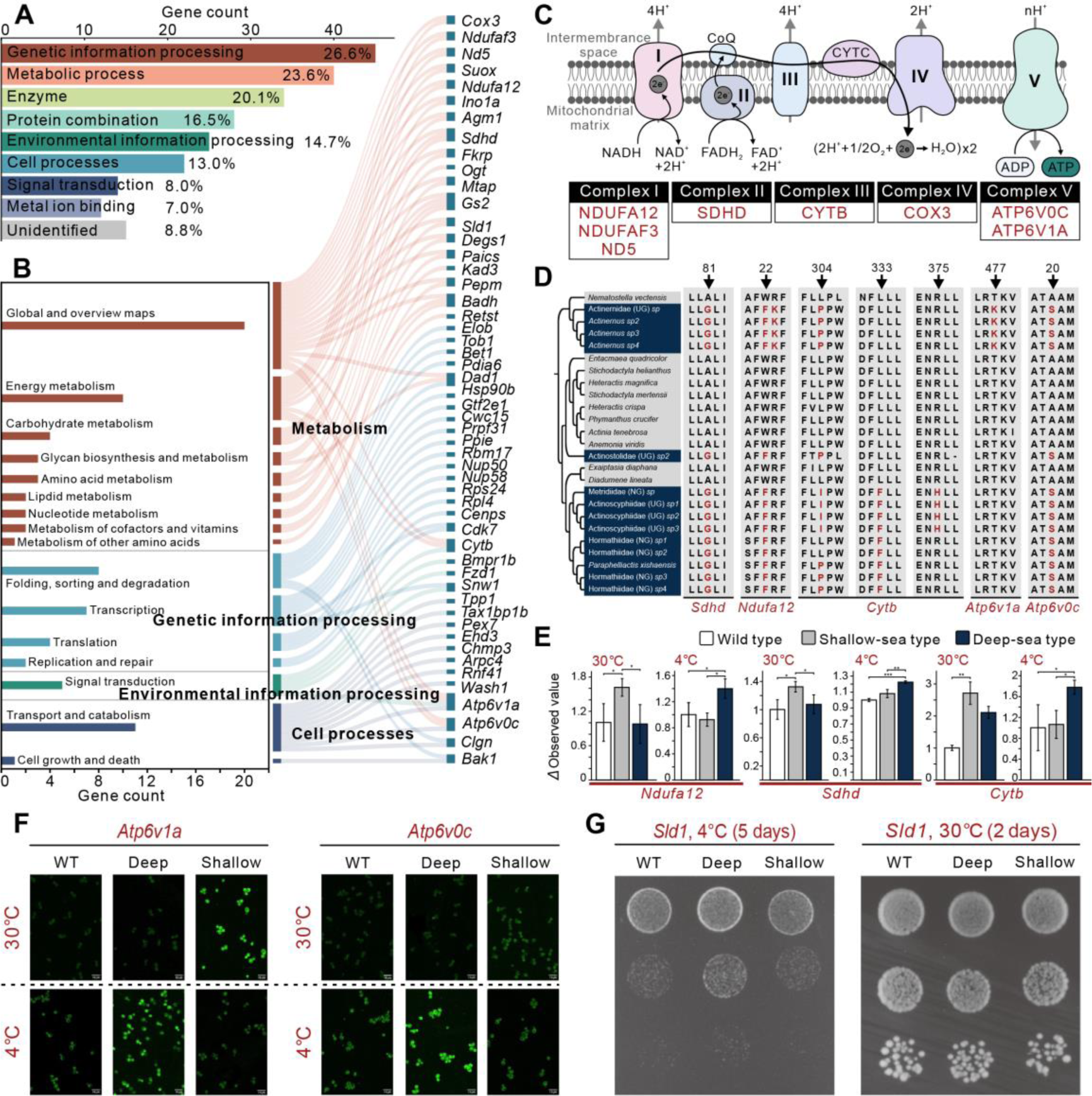
Functional analyses of deep-sea specific genes (DSGs). **(A and B)** Functional classification of 169 DSGs based on GO annotations (A) and KEGG annotations (B). Bar charts represent the number of genes in each category. **(C)** Schematic diagram of the electron transport chain, highlighting DSGs (in red) within each complex. **(D)** The mutation sites of five DSGs validated for functionality. Amino acid substitutions specific to deep-sea sea anemones are marked in red. **(E)** Functional activity validation of *Ndufa12*, *Sdhd*, and *Cytb* genes under 4°C and 30°C conditions. The analyses were performed using *Saccharomyces cerevisiae* (yeast). Mitochondria were extracted from yeast, and the activity of the corresponding genes was tested *in vitro*. **(F)** Functional activity validation of different genotypes of *Atp6v1a* and *Atp6v0c* under 4°C and 30°C conditions using the yeast experimental system. The fluorescence in the figure comes from LysoSensor Green DND-189 dye. Brighter fluorescence intensity indicates higher enzyme activity. **(G)** Growth of yeast strains carrying deep-sea and shallow-water variants of the *Sld1* gene, as well as wild-type yeast, under 4°C and 30°C conditions.

To functionally validate the adaptive significance of these deep-sea specific variants, we experimentally assessed enzyme kinetics for five key respiratory chain DSGs (Fig. 5D). For each gene, enzymes carrying the shallow-water-type amino acid variants exhibited higher activity at 30°C, whereas enzymes with deep-sea-specific amino acid variants consistently performed better at 4°C (Fig. 5E-F; raw data in Supplementary Table 13). This provides compelling functional evidence that these specific amino acid changes are critical determinants of enzyme activity under deep-sea relevant temperatures (2–4°C). Given that high hydrostatic pressure often exerts effects on biomolecules analogous to those of low temperature ^73–76^, these results strongly suggest direct protein-level adaptation to the combined physico-chemical challenges of the deep sea.

Among the DSGs, we also identified two fatty acid desaturase genes, *Sld1* and *Degs1* (Fig. 5B; Supplementary Figs. 14-15), responsible for introducing double bonds into fatty acids ^77–80^, a critical function for maintaining membrane fluidity in cold and high-pressure environments ^1,19,74,81^. We functionally assessed the *Sld1* variants using a Yeast spot assay. Yeast strains carrying the deep-sea-type *Sld1* exhibited significantly enhanced growth at 4°C compared to wild-type yeast or those carrying the shallow-water-type *Sld1* (Fig. 5G). This further validates the adaptive benefit conferred by deep-sea specific amino acid substitutions in optimizing protein function.

Collectively, the identification of numerous DSGs across diverse functional pathways signifies widespread protein tuning as a key adaptive strategy in deep-sea anemones. These findings also affirm the efficacy of our analytical approach in pinpointing functionally relevant amino acid substitutions across deep evolutionary divergences where traditional methods fail.

#### Functional compensation via gene family expansion

Beyond optimizing existing proteins and jettisoning non-essential functions, deep-sea anemones also employ gene family expansion as an adaptive mechanism. We identified 42 gene families significantly expanded in deep-sea lineages compared to their shallow-water counterparts, encompassing functions related to neurotransmission, membranes, DNA repair, etc (Fig. 6A). Notably, these expansions often result in total transcript levels for these functions in deep-sea species that are comparable to, or even exceed, those observed in shallow-water counterparts (Fig. 6A; Supplementary Fig. 16), suggesting a compensatory role. Intriguingly, the expanded neuronal gene families are heavily biased towards components associated with excitatory signaling (Fig. 6A). This includes adrenergic receptors (e.g., *Adrb1*, *Adrb2*), dopamine receptors (e.g., *Drd4*), muscarinic acetylcholine receptors (e.g., *Machr*), nicotinic acetylcholine receptors (e.g., *Chrna10*, *Chrnb2*), neuropeptide receptors (e.g., *Npffr2*, *Sifar*), kynurenine synthesis involved in neurotransmitter modulation (e.g., *Ido2*), and neurotransmitter transporters (e.g., *Slc6a11*, *Slc32a1*, and *Slc5a7*). This pronounced expansion of excitatory signaling components may represent an adaptive counterbalance to the inherently ‘suppressive’ nature of the extreme deep-sea environment on neuronal activity and overall metabolic rates.

**Fig. 6.**
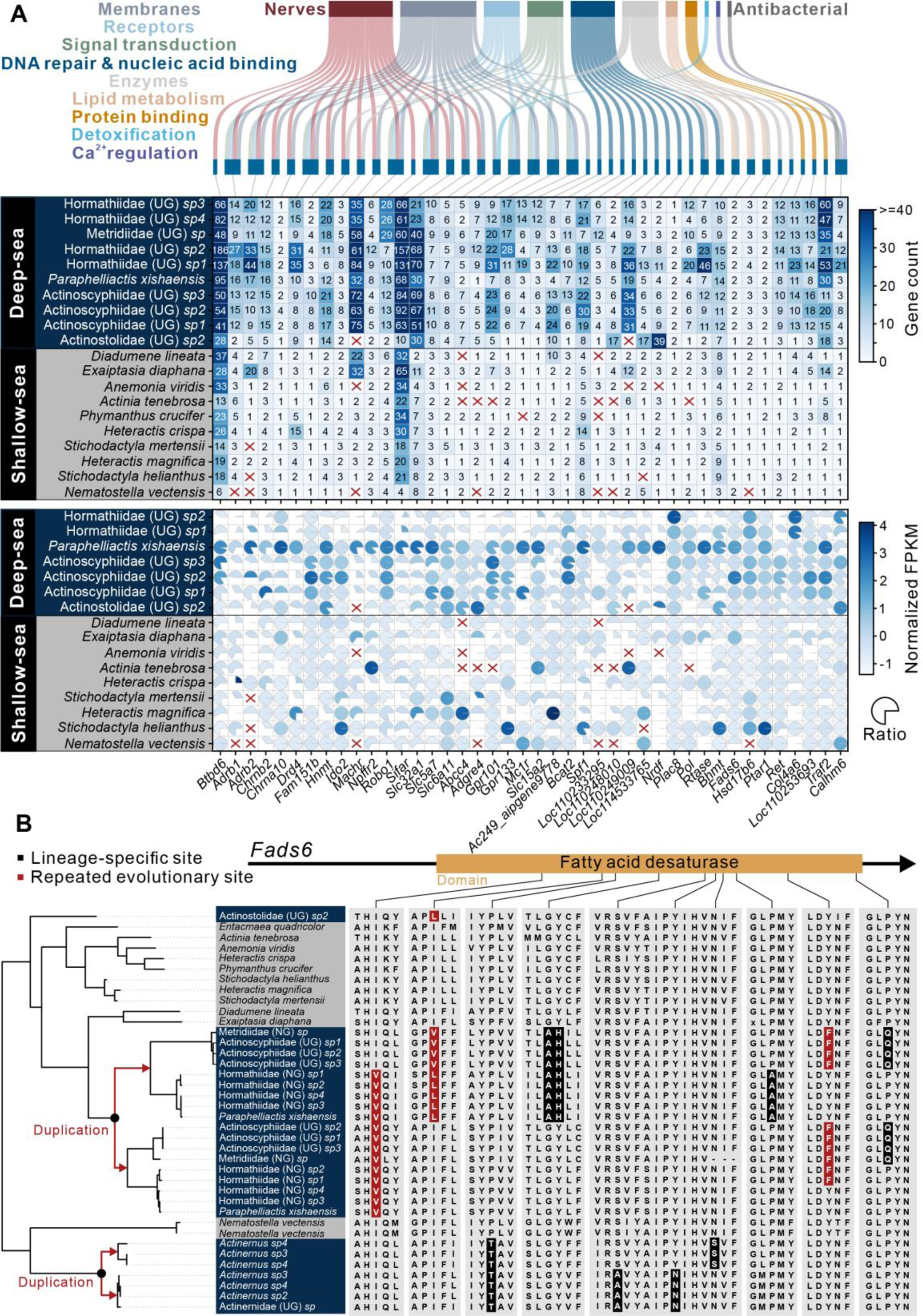
Repeated evolution of gene expansion in deep-sea anemones. **(A)** The Sankey diagram at the top illustrates the functional classification of expanded genes. The middle section shows the copy numbers of expanded genes in deep-sea and shallow-water anemones. Gene numbers are displayed using heatmap colors and are labeled in the figure. “X” indicates a gene number of 0. The bottom section shows the expression of expanded genes. The pie charts represent the proportion of transcribed gene copies relative to the total copy number. If all copies are expressed, the pie chart has a 360° arc. The color of the pie chart represents normalized expression levels (FPKM). **(B)** The duplication and amino acid variations of the fatty acid desaturase gene *Fads6*.

Furthermore, reinforcing the critical importance of lipid metabolism, the fatty acid desaturase *Fads6* not only underwent recurrent gene duplication events across different deep-sea lineages but also accumulated deep-sea specific amino acid variants and exhibited signatures of convergent amino acid evolution (Fig. 6B). This highlights how multiple adaptive mechanisms – gene duplication and protein sequence evolution – can converge on the same pathway to meet the physiological demands of deep-sea life.

### Macroevolutionary patterns of actiniarian diversification across depth

To investigate the macroevolutionary patterns of sea anemone diversification across different oceanic depths, we constructed a high-resolution phylogeny incorporating 267 actiniarian species (Supplementary Table 14), using the well-supported species tree (Fig. 2A) as a backbone constraint (Fig. 7A). This expanded phylogeny revealed an even more pronounced nested structure of deep-sea and shallow-water clades, underscoring the frequency of habitat transitions between these two major marine realms throughout actiniarian history. This tree also highlighted a tendency for extant deep-sea species to occupy relatively older phylogenetic branches (Fig. 7A).

**Fig. 7.**
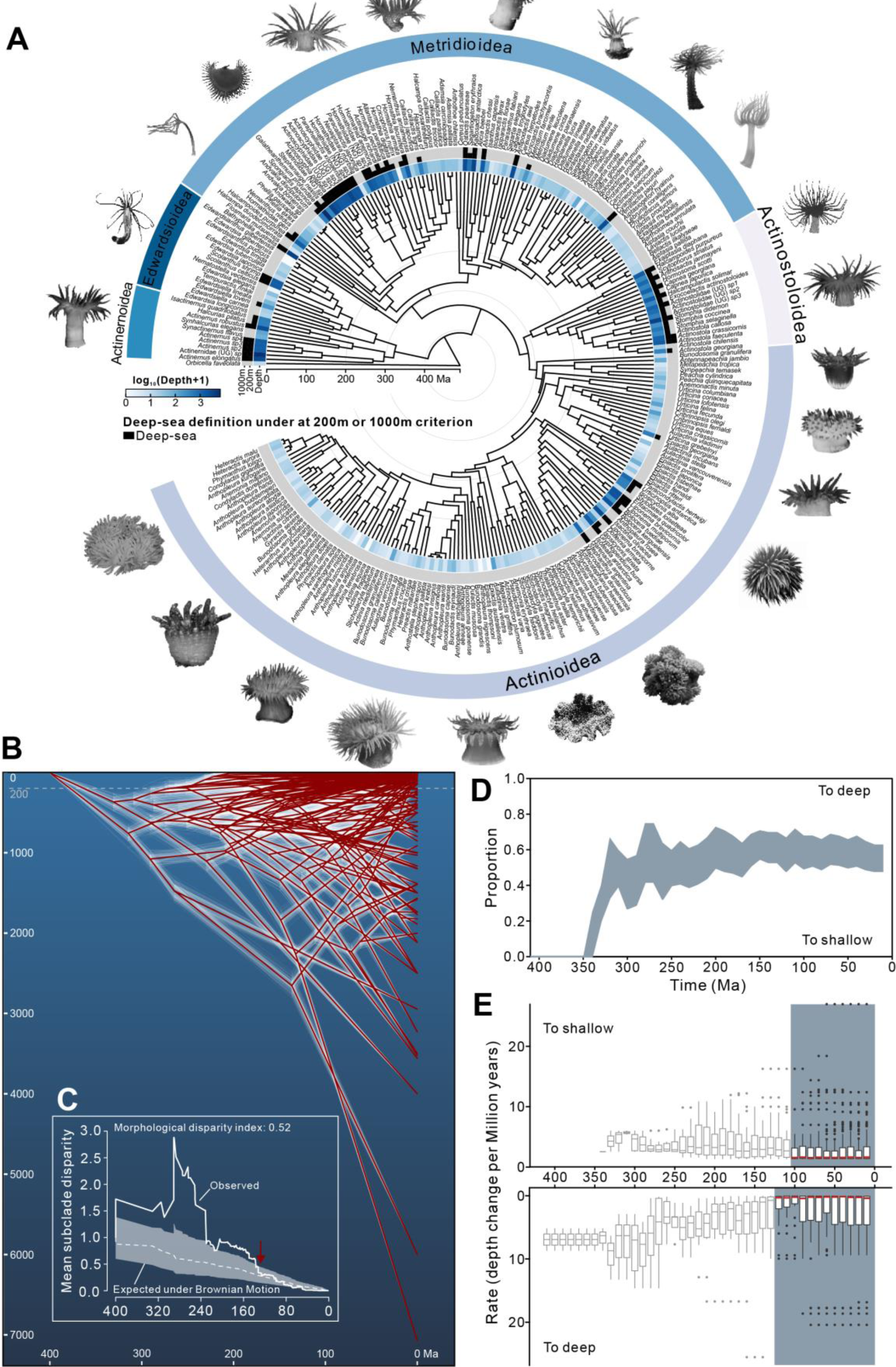
Macroevolutionary analyses of sea anemones across depth. **(A)** Phylogenetic tree of 267 sea anemones. Shades of blue represent species’ habitat depth, with darker tones indicating greater depth. Black blocks denote deep-sea taxa based on 200 m and 1,000 m depth criterion, respectively. **(B)** Ancestral depth reconstruction of sea anemones using the BMVT model. The y-axis indicates habitat depth (m), and the x-axis represents divergence time (Ma). White lines show results from 900 sampling replicates; the red line represents the mean depth across replicates. **(C)** Disparity-through-time (DTT) analysis of sea anemone habitat depth. The grey area denotes the 95% confidence interval of depth disparity from 10,000 Brownian motion simulations; the dashed line indicates the median of simulated values. The red arrow marks the end of significant deviation between observed and simulated values. **(D and E)** Evolutionary trend analyses of sea anemone depth transitions, using 10 Ma non-overlapping windows. (D) shows the 95% confidence interval of the proportion of transitions toward deeper or shallower habitats across 900 simulations. (E) illustrates the evolutionary rates of depth transitions toward deeper or shallower environments; the red line indicates the mean value.

To further dissect the depth-temporal dynamics of anemone evolution, we performed ancestral state reconstruction of habitat depth across the phylogeny. The results suggest substantial jumps in both time and depth between early nodes, a pattern corroborated by branch length distributions that strongly implies significant historical extinction or sampling gaps among early-diverging lineages (Fig. 7B, Supplementary Fig. 17). A disparity-through-time (DTT) analysis ^82^, quantifying the variance in habitat depth among contemporaneous lineages relative to the variance within subclades, revealed that observed disparity significantly exceeded that expected under a null model of Brownian motion evolution for a prolonged period, from the estimated crown age (∼434 Ma) until approximately 140 Ma (Fig. 7C). Such elevated early disparity could arise from either intense early convergent evolution or, more likely, an artifact created by substantial, biased extinction events.

The evaluation of the evolutionary direction of sea anemones revealed that, after approximately 150 Ma, the proportion curves of colonization into deep and shallow waters stabilized, with the frequencies of the two directions becoming roughly equal and remaining constant overall (Fig. 7D). Similarly, analyses of the colonization rates into deep and shallow waters indicated that the overall rates stabilized around 130 Ma (Fig. 7E). While certain species demonstrated the capacity for rapid, large-magnitude shifts in depth habitat, the average rates of colonization into deeper versus shallower waters appear remarkably balanced and constant over the last ∼130 million years. This conclusion was robust across three different metrics used to quantify evolutionary rates (Fig. 7E; Supplementary Fig. 18).

In summary, over the last ∼130 million years, sea anemones have stably migrated between deeper and shallower regions at nearly identical frequencies and rates (Fig. 7D-E). To further explore habitat transitions, we used SIMMAP ^83^ to reconstruct the history of transitions between discrete states defined as deep (≥200 m) and shallow (<200 m). The analysis overwhelmingly favored the All-Rates-Different (ARD) model allowing asymmetric rates (Supplementary Table 15). Paradoxically, the ancestral state reconstruction under this best-fit model inferred a deep-sea origin for Actiniaria (Supplementary Fig. 19), directly contradicting the shallow-water origin strongly supported by our independent convergent gene loss analysis (Fig. 3). This apparent conflict is readily reconciled by the extensive loss of early shallow-water lineages inferred from our other analyses (Fig. 7B-C; Supplementary Fig. 17). Such biased extinction would mislead standard phylogenetic ancestral state reconstruction methods, creating an artifactual signal for a deep-sea origin, while simultaneously explaining the observation that many extant deep-sea species reside on older phylogenetic branches. The convergent gene loss analysis (Fig. 3), being less susceptible to such sampling biases, likely provides a more accurate picture of the group’s ultimate shallow-water origins.

## CONCLUSIONS

By integrating genomic analyses of 15 deep-sea sea anemone species with phylogenomic and comparative genomic approaches, this study systematically reveals the evolutionary patterns of Actiniaria from shallow waters into the deep sea and delineates the multi-faceted adaptive strategies underpinning this transition:

First, employing a new analytical framework based on convergent gene loss, we provide compelling evidence that sea anemones originated in shallow waters and subsequently undertook multiple, independent invasions of the deep sea. This finding challenges the assumption that deep-sea lineages are ancient relics and offers a new paradigm for investigating the origins of deep-sea fauna. Intriguingly, while extant deep-sea anemone species often occupy older phylogenetic branches, our analyses suggest this is likely an artifact of extensive extinction among early shallow-water lineages – a clear case of “survivor bias”. Furthermore, we find that the overall efficiency of migration of sea anemones towards deeper and shallower habitats has been remarkably symmetrical over the last ∼130 million years. This underscores the need for caution when reconstructing ancestral states solely from extant taxa and highlights the pervasive influence of extinction history on perceived macroevolutionary patterns.

Second, our research uncovers the diverse molecular mechanisms facilitating sea anemone adaptation to the formidable deep-sea environment. At the genomic level, we document significant functional remodeling characterized by the systematic loss of genes associated with circadian rhythms, sensory perception (particularly light and temperature fluctuation), and photoprotection, alongside profound shifts in nutritional and reproductive strategies. These losses represent adaptive streamlining in response to the darkness, thermal stability, and energy limitations of the deep sea. Concurrently, at the gene level, specific amino acid substitutions optimize the function of key proteins, such as those involved in energy metabolism, enhancing their activity under high-pressure, low-temperature conditions. Complementing these changes, selective expansions of gene families related to neurotransmission and membrane function likely provide crucial compensatory mechanisms to counteract the suppressive effects of the deep-sea environment. This balanced strategy – shedding costly or unnecessary functions via gene loss while reinforcing essential processes through protein optimization and selective gene expansion – may represent a general paradigm for adaptation to extreme environments.

In conclusion, this study significantly advances our understanding of the evolution and adaptation of a key deep-sea metazoan group. Moreover, it provides a valuable conceptual framework and methodological reference point for future investigations into the adaptation of diverse life forms to Earth’s largest, yet least explored, biome.

## MATERIALS AND METHODS

### Sample Information

Fifteen deep-sea anemone specimens were collected between 2018 and 2019 during scientific expeditions utilizing the manned submersibles *Shenhai Yongshi* and *Jiaolong*. These specimens were retrieved from depths ranging from 432 to 6,000 meters in the Xisha Trough of the South China Sea and the Yap Trench, including its adjacent waters (Supplementary Table 1). Robotic arms were employed for precise and careful collection, ensuring the specimens remained in pristine condition. Upon retrieval to the research vessel, specimens were rinsed with 2 μm-filtered seawater to remove surface sediment, photographed, and labeled. They were subsequently preserved either at −80°C or in ethanol for downstream sequencing analyses. All specimens are curated by the Deep-Sea Science and Engineering Research Institute, Chinese Academy of Sciences, and are securely stored in the institute’s specimen repository.

### Genome sequencing and assembly

Genomic DNA was extracted from all deep-sea anemone specimens using the QIAGEN® Genomic DNA Extraction Kit (Cat#13323, Qiagen) according to the manufacturer’s standard protocol. The purity and quality of the extracted DNA were evaluated using a NanoDrop™ One UV-Vis spectrophotometer (Thermo Fisher Scientific, USA) and 1% agarose gel electrophoresis, respectively. DNA concentrations were precisely quantified with a Qubit® 3.0 Fluorometer (Invitrogen, USA). Based on DNA quality assessments, Oxford Nanopore sequencing was performed on four deep-sea sea anemone specimens: Actinostolidae (Unidentified genus, UG) *sp2*, Hormathiidae (NG) *sp1*, Hormathiidae (NG) *sp2*, and *Actinernus sp3*, using the PromethION platforms. In parallel, PacBio® HiFi sequencing was conducted on three additional specimens: Actinoscyphiidae (UG) *sp1*, Actinoscyphiidae (UG) *sp2*, and Actinoscyphiidae (UG) *sp3*. Additionally, all specimens were subjected to Illumina sequencing on the NovaSeq 6000 platform to generate 150 bp paired-end reads. Illumina reads were filtered using fastp v0.19.6 ^84^ with default parameters before downstream analyses.

The genome sizes of the 15 sea anemone specimens were first estimated based on quality-controlled Illumina reads using SOAPec v2.01 ^85^. For genomes sequenced via Oxford Nanopore technology, *de novo* assembly was performed using wtdbg2 v2.4.1 ^86^ with default parameters, followed by three rounds of polishing with Nextpolish v1.3.1 ^87^ to generate genome drafts. Genomes obtained through PacBio HiFi sequencing were assembled with hifiasm v0.19.8 ^88^ under standard parameters. Potential haplotype redundancies in the genome drafts were resolved using Purge_dups v1.0.1 ^89^ with default settings. For specimens with only Illumina paired-end reads, genome drafts were assembled using Platanus v1.2.4 ^90^ with default parameters, and scaffold continuity was improved where applicable using RagTag v2.1.0 ^91^ with its scaffold algorithm. The completeness of all genome assemblies was assessed using BUSCO v5.4.3 with the “metazoa_odb10” database ^92^.

The mitochondrial genomes of all sea anemone specimens were assembled and annotated using MitoFinder v1.4.1 ^93^ with the parameter “-o 5” based on quality-controlled Illumina paired-end reads. Additionally, the web service MITOS v2 ^94^ was employed to provide supplementary annotations for the mitochondrial genomes.

### Transcriptome sequencing and assembly

We collected tissues representing as many sea anemone cell types as possible (e.g., tentacle, column muscle, gonad, mesenterial filament, etc.). Total RNA was extracted from each tissue using TRIzol (Thermo Fisher Scientific, USA) and then pooled in equal amounts. After quality control of the RNA, libraries were constructed using the NEBNext Ultra RNA Library Prep Kit for Illumina (NEB, USA) and sent to Nextomics (Wuhan, China) for sequencing on the Illumina HiSeq 2000 platform. On average, 6 Gb of 150 bp paired-end reads were generated for each species.

All RNA reads were quality-controlled using fastp v0.19.6 ^84^ with default parameters. Transcriptome assemblies were generated using rnaSPAdes v3.12.0 ^95^ with standard parameters. Subsequently, open reading frames (ORFs) were predicted from all assembled transcripts using TransDecoder v5.5.0 (https://github.com/TransDecoder/TransDecoder).

### Genome annotation

Repetitive elements were identified using a combination of *de novo* and homology-based approaches. First, *de novo* libraries were constructed for each species using RepeatModeler v2.0.1 ^96^, and repetitive sequences were identified with RepeatMasker v4.0.7 ^97^. For the homology-based approach, repetitive elements were predicted by searching against Repbase using both RepeatMasker v4.0.7 ^97^ and RepeatProteinMask v1.36 ^98,99^. Tandem repeats were identified using Tandem Repeats Finder v4.07 ^100^ with the parameters “2 7 7 80 10 50 500 -d -h -ngs”.

Gene model annotation refer to previous studies^74,81,101,102^ using a combination of *ab initio* prediction, homology-based protein prediction, and transcriptome-based prediction methods: First, *ab initio* gene predictions were performed for the deep-sea sea anemones using Augustus v3.2.1 ^103^, with the gene training set of *Nematostella vectensis* (GCF_000209225.1). Subsequently, genome information for seven Anthozoa species (*N. vectensis* [GCF_000209225.1], *Actinia tenebrosa* [GCA_009602425.1], *Exaiptasia diaphana* [GCF_001417965.1], *Actinostola sp. cb2023* [GCA_033675265.1], *Acropora digitifera* [GCF_000222465.1], *Orbicella faveolata* [GCF_002042975.1], and *Stylophora pistillata* [GCF_002571385.1]) was retrieved from NCBI. The protein sets of these five species were mapped to the deep-sea anemone genomes using BLAT v. 35 ^104^ with default parameters, and gene models were predicted using GeneWise v2.4.1 ^105^ with default parameters. Finally, transcripts with complete ORFs were mapped to the deep-sea sea anemone genomes using BLAT v. 35 ^104^ and further processed using PASA (Program to Assemble Spliced Alignments) v2.5.2 ^106^ to refine gene structures. All prediction results were integrated using EvidenceModeler v1.1.1 ^107^ with the following weighting scheme: TRANSCRIPT:10, PROTEIN:10, and ABINITIO PREDICTION:1, resulting in the final non-redundant gene set. Functional annotation of all gene models was performed with InterProScan v5.39-77.0 ^108^.

### Phylogenetic analyses

To investigate the phylogenetic positions of deep-sea and shallow-water anemones, we retrieved genome and transcriptome data for 26 published sea anemones from public databases (Supplementary Table 6). Combined with the 15 deep-sea anemones analyzed in this study, a total of 41 species were included, covering all known superfamilies within Actiniaria. *Orbicella faveolata*, which belongs to Scleractinia, was used as the outgroup for phylogenetic analysis.

Based on the whole-genome dataset from the aforementioned species, 1,849 orthologous gene sets were identified through Reciprocal Best Hit (RBH) analysis. These orthologous gene sets’ CDS sequences were aligned using MAFFT v7.407 ^109^ with default parameters, and low-quality alignment regions were trimmed with trimAl v1.5.0 ^110^ under parameter “-automated1”. Phylogenetic trees were constructed using IQ-TREE v2.2.2.4 ^111^ with the parameter settings “-m MFP-alrt 1000” for the trimmed sequences, including both gene trees of each molecular marker and a tree of all concatenated markers. Additionally, species tree analysis was performed using ASTRAL v5.5.9 ^112^ with default parameters.

For mitochondrial genes, 13 protein-coding genes’ CDS sequences were concatenated into a supergene, and phylogenetic analysis was conducted using the same method described above for gene tree construction.

Divergence times for sea anemones were estimated using the MCMCtree program in the PAML package v4.9h ^113^, based on three time calibration points: the most recent common ancestor (MRCA) of *Metridium senile* and *Anemonia viridis* (estimated at ∼369–504 Ma), the MRCA of *Anemonia viridis* and *Aiptasia* (estimated at ∼334–339 Ma), and the MRCA of *Orbicella faveolata* and *Metridium senile* (estimated at ∼437–539 Ma). These calibration times were obtained from the website TimeTree and confirmed through additional studies ^17,19^.

### Quartet Sampling

Based on the species tree, a Quartet Sampling (QS) analysis was performed on all gene trees involved in the ASTRAL v5.5.9 ^112^ analysis using the quartet_sampling.py script (https://www.github.com/fephyfofum/quartetsampling) ^114^ to evaluate the consistency and branch support of each node in the species tree. The parameter settings were as follows: the number of repetitions was set to 1,000 (-reps 1,000), the log-likelihood threshold was kept at the default value (-lnlike 2), and the minimum overlap of sampled loci for all taxa in a quartet was set to 20,000 (-min-overlap 20,000). Using the resampled quartet counts, we calculated the following metrics for each internal branch of the focal tree: Quartet Concordance (QC; consistency of quartet information), Quartet Differential (QD; presence of secondary evolutionary histories), Quartet Informativeness (QI; informativeness of the quartet), and Quartet Fidelity (QF; reliability of individual taxa in the tree). These metrics were used to assess the confidence, consistency, informativeness of internal nodes, and the reliability of each terminal branch.

### Dispute genealogy diagnoses

This study carefully addressed all conflicting topologies, which may have arisen due to data selection, algorithmic differences, or other biological processes introducing phylogenetic discordance.

For controversial branches in the phylogenetic tree, we used the DiscoVista v1.0 ^115^ software to evaluate topological bias and complexity. The specific procedure was as follows: First, we set the parameters “-k1 -m 5” to count the quartet frequencies of disputed branches. Next, in the support assessment of conflicting topologies, we considered 14 monophyly hypotheses (taking into account all discordant topologies between gene trees and species tree, as well as between species tree and previously published phylogenetic trees), including: (1) Actinostoloidea; (2) Actinernoidea; (3) Actinernidae (UG) *sp*/*Actinernus sp2*/*Actinernus sp3*/*Actinernus sp4*; (4) Metridioidea; (5) Actinostoloidea/Actinioidea/Metridioidea; (6) Actinioidea; (7) Edwardsioidea; (8) Edwardsioidea/Actinernoidea; (9) *Actinernus sp2*/*Actinernus sp4*/*Actinernus sp3*; (10) Actinostoloidea/Actinioidea; (11) *Actinernus sp4*/*Actinernus sp3*; (12) *Actinernus sp3*/*Actinernus sp4*/Actinernidae (UG) *sp*; (13) Actinioidea/Metridioidea; (14) *Actinernus sp4*/Actinernidae (UG) *sp*. Support exceeding 75% was considered strong.

In addition, this study employed the concordance factors algorithm in IQ-TREE v2.2.2.4 ^111^ to assess the support for gene sets within the phylogenetic tree topology ^116^. The analysis was conducted with the following parameter settings: -lmap ALL -m GTR+F+R5 -n 0. Among these, GTR+F+R5 was determined as the best-fit model by IQ-TREE v2.2.2.4 ^111^.

Gene flow and incomplete lineage sorting (ILS) are two major factors contributing to phylogenetic conflicts ^117–120^. To comprehensively evaluate the impact of these two factors on the phylogeny inferred in this study, we used the T1 model (representing ILS) in the MSCquartets ^121^ algorithm to examine all gene trees in the dataset. Specifically, the gene trees of the 1,849 orthologous gene sets were used as input data, and the quartet frequencies (quartet concordance consistency factors, qcCF) for each node in these trees were calculated using MSCquartets ^121^. Under the T1 model, the function quartetTreeTest was applied to test all quartet counts at four different rejection thresholds (α): 0.01, 0.001, 1e-04, and 1e-06.

### Demographic history

This study used the PSMC model ^122^ to estimate the population history dynamics of deep-sea and shallow-water anemones. First, heterozygous sites were extracted from BAM files generated by mapping NGS reads, using SAMtools v1.10-76-g65c8721 ^123^ with the parameters “mpileup -q 20 -Q 20”. Subsequently, the base substitution rate of the species was estimated using the penalized likelihood method implemented in r8s v1.81^124^, based on the divergence time calibration and branch lengths from the species tree topology (Supplementary Table 8). Finally, the PSMC analysis was conducted with the following parameters: -t 15 -r 5 -p “4+25*2+4+6”. The generation time for all species was set to 1 year ^17,19^.

### The repeated loss of gene families

This study proposed a suitable workflow to identify gene families that are commonly lost among multiple lineages, tailored to the characteristics of the data. Detailed protocols are shown in Fig. 3B.

#### Step 1: Gene group

This study selected the protein dataset of the following sea anemone species: Actinoscyphiidae (UG) *sp1*, Actinoscyphiidae (UG) *sp2*, Actinoscyphiidae (UG) *sp3*, Hormathiidae (NG) *sp1*, Hormathiidae (NG) *sp2*, Actinostolidae (UG) *sp2*, *Actinernus sp3*, *Actinia tenebrosa*, *Exaiptasia diaphana*, *Paraphelliactis xishaensis*, *Nematostella vectensis*, *Stichodactyla mertensii*, *Anemonia viridis*, *Heteractis crispa*, *Stichodactyla helianthus*, *Diadumene lineata*, *Heteractis magnifica*, *Phymanthus crucifer*, and *Entacmaea quadricolor*. Gene family clustering analysis was conducted using OrthoFinder v2.3.1 ^125^ with the following parameters: -M msa -T iqtree -s species.tre - y -S diamond, where “species.tre” represents the species tree topology of these taxa.

The analysis ultimately identified 21,791 N0-level orthogroups.

#### Step 2: Initial filtering

To preserve as many potentially lost gene families as possible, relatively lenient filtering criteria were applied in this step. The specific conditions were adjusted based on the analysis, as detailed in Fig. 3B. This process resulted in the identification of candidate lost orthogroups.

#### Step 3: Gene check in all species

In this step, the protein sequences of candidate orthogroups were used as queries to manually identify potential orthologs in the genomes of all species, using a combination of BLAT v. 35 ^104^ and GeneWise v2.4.1 ^105^. Results with lengths less than 30% of the query were filtered out. Subsequently, all potential orthologs were further confirmed for homology using RAxML v8.2.13 ^126^ in combination with OrthoFinder v2.3.1 ^125^. All software analyses were conducted with default parameters.

#### Step 4: Further filtering

To prevent some results from being mistakenly excluded due to poor genome assembly quality, relatively lenient filtering criteria were again applied in this step. The specific conditions were adjusted based on the analysis, as detailed in Fig. 3B. This step preliminarily identified the lost orthologs.

#### Step 5: Final inspection and filtering

This step relied on transcriptomes and NGS reads to further identify missing or retained genes that may have been overlooked due to poor genome assembly quality (in six deep-sea species). A similar approach can be found in Xu *et al.* ^81^ and Wang *et al.* ^74^.

For transcriptome analysis: Transcripts with complete ORFs were analyzed using TransDecoder v5.5.0 (https://github.com/TransDecoder/TransDecoder) to generate a protein set. The orthologs identified in the previous step were used as queries to search the protein set with Diamond v0.9.24.125 ^127^ in combination with RAxML v8.2.13 ^126^ to evaluate the loss or retention of genes within species.

For NGS reads analysis: Conserved regions of the identified orthologs were determined, and their positions in the gene-retained genomes were annotated based on GFF files. Subsequently, NGS reads from the six deep-sea species were mapped to these gene-retained genomes using BWA v0.7.12-r1039 (https://github.com/lh3/bwa)^128^. The presence or absence of a gene in a species was assessed by examining the read coverage in the conserved regions.

Finally, the results of both analyses were integrated and filtered based on specific parameters in Fig. 3B to obtain the final set of orthologs

#### Step 6: Robust loss checking

The final orthologs were used as queries to manually examine the genomes of all 25 species using a combination of BLAT v. 35 ^104^ and GeneWise v2.4.1 ^105^. This process assessed whether functionally similar paralogs were retained. For orthologs where no similar paralogs were identified, we considered the corresponding function to be completely lost, and these genes were designated as robust loss genes.

### Investigation of symbionts in sea anemones

Based on the previous reports ^47–49^, we retrieved the most comprehensive database of symbiotic algae associated with cnidarians to date. From this database, we extracted all ITS2 sequences classified as Clade A–I endosymbionts. Using BWA v0.7.12-r1039 (https://github.com/lh3/bwa) ^128^, NGS reads from both deep-sea and shallow-water anemones were mapped to the ITS2 database. Finally, the number of successfully mapped reads was quantified to evaluate whether the sea anemones harbored symbiotic algae.

### Histological staining and *in situ* hybridization

To compare the histological characteristics of gonads between deep-sea and shallow-water anemones, this study selected gonadal samples from shallow-water species *Exaiptasia diaphana* and *Actinia tenebrosa*, as well as deep-sea species Actinoscyphiidae (UG) *sp1*, Hormathiidae (NG) *sp1*, Hormathiidae (NG) *sp3*, and *Actinernus sp4*, based on sample availability. For shallow-water anemones, conventional paraffin sectioning techniques were used for histological observation.

Since deep-sea anemones were preserved as frozen samples, cryosectioning techniques were employed to maximize sample integrity. The preparation of sections and hematoxylin-eosin (HE) staining followed previously published protocols ^129^ and were appropriately optimized based on the preservation conditions of the samples (e.g., sample fixation, section thickness, etc.).

For the *in situ* hybridization experiment, tissue samples of *Exaiptasia diaphana* were rinsed in PBS and then fixed in *in situ* hybridization fixative at 4°C for 12 hours. Subsequently, cubic tissue blocks (approximately 1 cm per side) were dissected from the columnar region containing the gonads. The tissue blocks were dehydrated sequentially in 15% and 30% sucrose solutions.

The dehydrated tissues were cryosectioned longitudinally, and the sections were air-dried before being fixed in 4% paraformaldehyde for 10 minutes. The sections were washed three times with PBS (pH 7.4) for 5 minutes each. After air cooling, proteinase K (20 μg/mL) was added, and the sections were digested at 40°C, followed by three washes in PBS.

Pre-hybridization was performed at 40°C for 1 hour, followed by probe hybridization at 40°C overnight. After hybridization, the sections were washed sequentially with 2× SSC, 1× SSC, and 0.5× SSC. Next, 60 μL of pre-heated branched probe hybridization solution was applied, and the sections were hybridized at 40°C for 45 minutes, followed by washing. Finally, a signal probe hybridization solution (1:200 dilution) was added, and the sections were incubated at 40°C for 3 hours, with washing steps performed as described above.

After probe hybridization, the sections were counterstained with DAPI staining solution in the dark for 8 minutes to label nuclei. The sections were mounted and observed under a Nikon upright fluorescence microscope. Images were captured using the following excitation and emission wavelengths: DAPI (359 nm/457 nm, blue fluorescence); iF488-Tyramide (491 nm/516 nm, green fluorescence); iF546-Tyramide (541 nm/557 nm, red fluorescence).

### Deep-sea specific genes

In investigating amino acid-level adaptations in proteins, traditional methods based on selection pressure analyses to identify positive selection or rapid evolutionary events are not suitable for the long evolutionary span of sea anemones involved in this study. Therefore, we introduce an analysis pipeline based on information entropy to detect deep-sea lineage-specific amino acid variations in conserved regions of genes. The specific protocols are as follows:

Amino acid site alignment: We used MACSE v2.06 ^130^ software to perform codon alignment of the aforementioned orthologous genes in sea anemones. The parameters were set as “-prog alignSequences -gc_def 1”.

Assessment of amino acid site conservation: The conservation level of each amino acid site was evaluated using an information entropy-based algorithm. The formula is as follows:

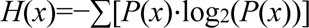

In the formula, *P*(*x*) represents the frequency of a specific amino acid occurring at a given site. A smaller information entropy value indicates that the site is more conserved. For sites where amino acid deletions (-) occur, if the deletion proportion exceeds 60%, the entropy value of the site is marked as “x”, and the site is excluded from subsequent analyses.

Detection of deep-sea-specific amino acid variation sites: Based on the aligned sequence file, we used an in-house script to identify amino acid variation sites specific to each deep-sea lineage (the four deep-sea lineages shown in Fig. 2A) compared to all shallow-water lineages. The criteria for detection were as follows: the test deep-sea lineage must share a common amino acid, while all shallow-water species must share another common amino acid.

Evaluation of upstream and downstream conservation of variation sites: The conservation of the regions surrounding the variation sites (within a range of 10 amino acids upstream and downstream) was assessed using the Theta-w value. The formula is as follows:

Theta-w = (AA value) / (counted AA number)

In the formula, the AA value represents the weighted sum of *H*(*x*) values for amino acid sites within the statistical region. The weighting rules are as follows:

- If *H*(*x*) < 0.01, the weight is 1.
- If 0.01 ≤ *H*(*x*) ≤ 0.5, the weight is 0.5.
- If 0.5 < *H*(*x*) ≤ 1, the weight is 0.3.

Filtering and integration: Finally, we retained results with Theta-w > 0.6. Based on these results, we further selected genes that exhibited specific amino acid variations in all deep-sea lineages. It is important to note that the specific amino acid variations were not required to occur at the same sites across different lineages. The genes identified through this process were defined as Deep-Sea Specific Genes (DSGs).

To understand the functional characteristics of DSGs and their potential roles in deep-sea adaptation, we first performed Gene Ontology (GO) annotation for all DSGs and categorized them according to functional categories. Subsequently, we used the KOBAS platform ^131^ (http://bioinfo.org/kobas/) to conduct Kyoto Encyclopedia of Genes and Genomes (KEGG) pathway annotation and enrichment analysis for these genes. Enrichment results with FDR-corrected *p*-values less than 0.05 were considered significant. Next, based on the pathway classification standards provided by the KEGG Pathway Database (https://www.kegg.jp/kegg/pathway.html), the annotated genes were categorized and statistically analyzed.

### Preparation of yeast expression strains and enzyme activity assay

#### Plasmid construction

In this study, yeast expression plasmids were constructed for both wild-type and mutant forms of *Ndufa12*, *Sdhd*, *Cytb*, *Atp6v1a*, and *Atp6v0c*, as well as *Sld1* (deep-type) and *Sld1* (shallow-type). *Ndufa12*, *Sdhd*, and *Atp6v0c* were derived from the cold-adapted Hormathiidae (NG) *sp2* (sample ID: HK2SQW55). The mutant forms included Phe22 mutated to Trp (*Ndufa12*), Gly81 mutated to Ala (*Sdhd*), and Ser20 mutated to Ala (*Atp6v0c*). *Cytb* was derived from the cold-adapted Actinoscyphiidae (UG) *sp3* (sample ID: HK2SQW43), with three mutations: Ile304Leu, Phe333Leu, and His375Arg. *Atp6v1a* was derived from the cold-adapted *Actinernus sp3* (sample ID: HK2SQW54) with one mutation, Lys477Thr. *Sld1* (deep-type) was derived from the cold-adapted Hormathiidae (NG) *sp2* (sample ID: HK2SQW55), while the cold-intolerant *Sld1* gene was derived from the cold-intolerant species *Diadumene lineata* (GenBank: GCA_918843875.1). All sea anemone samples, except for *Diadumene lineata*, are stored at the Institute of Deep-Sea Science and Engineering, Chinese Academy of Sciences. The DNA sequences of the wild-type genes were codon-optimized and synthesized by Sangon Biotech (Shanghai, China). These sequences were cloned into the pRS416 vector, flanked by the TEF1 promoter and CYC1 terminator. The mutant constructs were generated by introducing mutations into the wild-type sequences via PCR. All plasmids were further validated by sequencing.

#### Yeast transformation

A single fresh colony was inoculated into the appropriate medium and cultured overnight at 30°C. The culture was then diluted to an OD600 of 0.1 and transferred to 5 mL of fresh medium, followed by incubation at 30°C until the OD600 reached approximately 0.6. Cells were collected and washed with 1 mL sterile ddH_2_O and 500 μL of 0.1 M LiOAc, then resuspended in 100 μL of 0.1 M LiOAc. Next, 20 μL of the cell suspension was mixed with 80 μL of transformation buffer (containing 58.6 μL 50% PEG3350, 7.7 μL 1 M LiOAc, 9.0 μL DMSO, and 4.7 μL ssDNA) and the plasmid for gene expression. The mixture was gently pipetted to mix thoroughly and incubated at 30°C for 35 minutes, followed by a 15-minute heat shock at 42°C. The cells were centrifuged to remove the supernatant, resuspended in 200 μL sterile ddH_2_O, and spread onto the selective solid medium. The plates were incubated at 30°C for 3 days.

#### Respiratory chain complex activity assay

A fresh single colony was inoculated into 5 mL of liquid medium and cultured overnight at 30°C. The culture was then transferred to 100 mL of liquid medium and diluted to an OD600 of 0.1, followed by incubation at 30°C until the OD600 reached 0.8. Cells were then collected and resuspended in 1 mL of 1× PBS buffer (P1020, Solarbio), followed by centrifugation at 5000×g for 1 minute at room temperature. This process was repeated twice. Mitochondria were extracted using a yeast mitochondrial extraction kit (EX2900, Solarbio). The isolated mitochondria were disrupted by sonication at a power of 200 W, using 5-second pulses with 10-second intervals, repeated 15 times. Protein quantification of the respiratory chain complexes obtained from the lysed mitochondria was performed using a BCA protein assay kit (23225, Thermo Fisher). The activities of three electron transport chain complex proteins NDUFA12, SDHD, and CYTB were measured using microassay methods according to the manufacturer’s instructions. NDUFA12: Mitochondrial respiratory chain complex I activity assay kit (BC0515, Solarbio). SDHD: Mitochondrial respiratory chain complex II activity assay kit (BC3235, Solarbio). CYTB: Mitochondrial respiratory chain complex III activity assay kit (BC3245, Solarbio). Protein activity assays were conducted under two temperature conditions, 30°C and 4°C, as specified by the experimental design. Finally, enzymatic activities of mitochondrial electron transport chain complexes were measured at the corresponding temperature using a multifunctional microplate reader. Each group included three technical replicates. The complex activity values were normalized as follows: Relative enzymatic activity = measured value / mean value of three technical replicates of the wild type.

#### Vacuolar enzyme activity assay

A fresh single colony was inoculated into the appropriate medium and cultured overnight at 30°C. The culture was then diluted to an OD600 of 0.1 and transferred to 5 mL of fresh medium, followed by further incubation at 30°C until the OD600 reached approximately 0.8. Two tubes of 1 mL culture were collected and centrifuged to remove the supernatant. The cells were washed with 1 mL of sterile PBS buffer, centrifuged again to remove the supernatant, and this process was repeated twice. The cells were then resuspended in 800 μL of PBS buffer. Yeast cells were then stained using LysoSensor Green DND-189 (L7535, Thermo Fisher). A 4 μL aliquot of the dye was added to the resuspended cells, mixed thoroughly, and incubated separately at 4°C and 30°C for 6 hours. After incubation, the cells were centrifuged to remove the supernatant and washed three times with 1 mL of sterile PBS buffer. The cells were then resuspended in 1 mL of sterile PBS and immediately observed under a Nikon AXR laser scanning confocal microscope for fluorescence imaging. The green fluorescence protein (GFP) was excited using a 488 nm argon laser, and the emission spectrum was collected between 499 and 542 nm. Image data acquisition and quantitative analysis were performed using Fiji software to ensure image quality and reproducibility.

#### Yeast spot assay

A single fresh colony was inoculated into SC-URA liquid medium and cultured overnight at 30°C. The culture was then adjusted to an OD600 of 1.0. Yeast strains were either directly spotted or subjected to 10-fold serial dilutions prior to spotting onto SC-URA solid medium. The plates were incubated at 30°C and 4°C for several days, after which photographs were captured.

### Gene family expansion

This analysis involved 7 deep-sea anemones and 10 shallow-water anemones. Whole-genome proteins of all species were clustered using OrthoFinder v2.3.1 ^125^. Based on the clustering results at the N0 level, we scored the gene family expansions for each deep-sea anemone relative to the 10 shallow-water anemones using the Z-score.

Results with a Z-score > 1.96 were considered significantly expanded. Gene families that were significantly expanded in at least 6 out of the 7 deep-sea anemones were defined as deep-sea-expanded gene families. These results were subsequently validated in Metridiidae (UG) *sp*, Hormathiidae (NG) *sp3*, and Hormathiidae (NG) *sp4*. The validation method followed Step 3 in the “The repeated loss of gene families” section. Next, these expanded gene families were categorized based on their functions using GO and KEGG annotations.

### Transcriptome expression analysis

To evaluate gene transcription levels, this study used HISAT2 v2.2.1 ^132^ to align quality-controlled RNA-seq reads to the reference genome. Subsequently, based on the gene annotation information of the species, the StringTie v2.1.1 ^133^ software was used to calculate paired reads counts for each gene and compute fragments per kilobase of exon per million mapped reads (FPKM) for subsequent transcript quantification analysis.

In addition, this study also considered normalizing transcriptional values using internal controls. Based on previous studies ^134–139^, we retrieved 19 candidate genes that could serve as internal control genes, including 18S rRNA, 28S rRNA, 40S rRNA, 60S rRNA, and *Actin*, among others (Supplementary Table 16). Based on the expression stability of these genes across species (Supplementary Fig. 20), we selected 40S_3, 40S_4, and 40S_9 (gene symbols were represented in Supplementary Table 16) as internal controls for comparative transcriptomic analyses.

### Phylogeny of 267 sea anemones

To construct a higher-resolution phylogenetic tree of Actiniaria in order to better serve macroevolutionary analyses, this study retrieved all available Actiniaria data from public databases. After data cleaning, 267 species were retained for subsequent analyses (Supplementary Table 17). *Orbicella faveolata* was selected as the outgroup for phylogenetic analysis. Using the time-calibrated species tree obtained above as a backbone, this study inferred the phylogenetic relationships of 267 Actiniaria species based on concatenated sequences of 12S rRNA, 16S rRNA, *Cox3*, *Cox1*, 18S rRNA, and 28S rRNA. Phylogenetic analysis was conducted using IQ-TREE v2.2.2.4 ^111^ with the parameter “-m MFP -alrt 1000”. Divergence times for each node were subsequently estimated using the MCMCTree program in PAML v4.9h ^113^.

### Ancestral status assessment

To evaluate the evolutionary patterns of sea anemones in the “time-depth” dimension, we collected depth distribution data for the aforementioned 267 species. After verifying the distribution data, the median depth was calculated for each species and used as its depth parameter (Supplementary Table 14).

Based on the phylogenetic relationships and depth distribution data of these species, we used the fossilBM ^140^ program to infer the distribution states at all nodes in the evolutionary tree under the BMVT model. This algorithm is considered to better accommodate data missingness in large-scale phylogenetic studies ^140–142^. The analysis was conducted with 2,000,000 Markov chain Monte Carlo (MCMC) iterations, sampling every 2,000 iterations. The first 200,000 generations (i.e., the first 10%) were discarded as burn-in. The remaining samples were then aggregated for further analysis.

Finally, using the high-resolution time-calibrated tree of sea anemones, we conducted a disparity-through-time (DTT) analysis ^82^ in the R package *geiger* ^143^ to evaluate the evolutionary patterns of sea anemones along the “depth” axis.

Additionally, the analysis included 10,000 random simulations to assess the expected evolutionary trajectory of depth distribution under unconstrained conditions.

### Evolutionary trend assessment

First, we defined colonization events as follows: when the depth of the parent node is shallower than that of the child node, it is considered a shallow-to-deep colonization event; conversely, when the parent node’s depth is deeper than the child node’s, it is considered a deep-to-shallow colonization event. Colonization events are assumed to occur with a uniform probability over the time interval between the parent and child nodes. All evaluations were conducted within a 10 Ma non-overlapping sliding window (step size = 10 Ma).

Based on the 900 trees collected from the BMVT model, we calculated the proportion of “deep-to-shallow” and “shallow-to-deep” events within each sliding window for each tree. Finally, the results from all 900 trees were aggregated, and the 95% confidence intervals were calculated.

We also evaluated the mobility rates of “deep-to-shallow” and “shallow-to-deep” colonization events, specifically the average depth change per unit of time for sea anemones. Three statistical metrics were used in this study to reflect the evolutionary rates:

*R*=(D1-D2) / (|t1-t2|) ^144^

*V*=(ln(D1) - ln(D2)) / (|t1-t2|) ^144,145^

*H*=(ln(D1) - ln(D2)) / (S*|t1-t2|) ^144,146^

*R*: Evolutionary rate, calculated as the average depth change per unit time; *V*: Darwinian rate, defined as the rate of change in the natural logarithm of depth; *H*: Haldane rate, incorporating the variance of depth changes among sampled trees; D1: Average depth (m) of the parent node; D2: Average depth (m) of the child node; t1: Time (Ma) of the parent node; t2: Time (Ma) of the child node; S: Standard deviation of ln(D1) and ln(D2) across all 900 sampled trees for the same node.

### SIMMAP analysis

We defined the status of all sea anemones using 200 m as the boundary between deep and shallow seas ^1,19,74^. Subsequently, we used the *simmap* function (via the *ace* function in the R package) to evaluate the fit of three models: ER (equal-rates model), SYM (symmetrical model), and ARD (all-rates-different model) ^83^ and reconstructed the inhabitant state of all nodes.

## ACKNOWLEDGMENTS

We acknowledge the crews of the research vessel *Tansuo 1* and the pilots of the HOV *Shenhai yongshi*. We would like to express our sincere gratitude to Professor Zhiyong Yue from Xi’an International University for his assistance in experimental exploration. We are also indebted to our collaborators, Baosheng Wu, Huishan Yue, Xueli Gao, Zhenhong Pan, Wenshi Xue, and Chenyi Yang, for their instrumental assistance with bioinformatics and experimental analyses. This project was supported by the National Key R&D Program of China (2022YFC3400300, 2023YFC2809300, 2016YFC0304905), by the Fundamental Research Funds for the Central Universities, Northwestern Polytechnical University (D5000220464), the National Natural Science Foundation of China (32100367, 32370452), the 1000 Talent Project of Shaanxi Province to Kun Wang and Qiang Qiu, and Shaanxi Postdoctoral Research Project (2023BSHGZZHQYXMZZ53).

## AUTHOR CONTRIBUTIONS

C.F., H.Z., Q.Q., and K.W. conceived the study. H.Z. and Y.Z. collected the sea anemone. Y.Z. performed the morphological validation experiments of the sea anemone samples. P.X. performed the genome assembly and genome annotation. P.X., C.L., W.X., C.Z., M.H., Ye.L., Yu.L., J.Z., T.Q., Y.Y., and P.L. performed the data analysis. X.W., P.X., H.S., Y.Z., and L.Q. performed the yeast gene editing experiments. C.F., K.W., P.X., and X.W. wrote the manuscript. All authors provided comments and approved the manuscript for submission and publication.

## COMPETING INTERESTS

The authors declare no competing interests.

## DATA AVAILABILITY

All sequencing reads and genome assemblies have been deposited into CNGB Sequence Archive (CNSA, https://db.cngb.org/cnsa/) ^147^ with accession number CNP0007216. The CNSA sample accession numbers for all assembled genomes are documented in Supplementary Table 4. All the scripts and code of the analysis of this study are available at https://github.com/Peidong-Xin/deep-seaAnemone.

## Notes

### Competing Interest Statement

The authors have declared no competing interest.

## REFERENCES

1 Jamieson, A. The hadal zone: life in the deepest oceans. (Cambridge University Press, 2015).

2 Tyler, P. Ecosystems of the deep oceans. (Elsevier Science, 2003).

3 Higgs, N. D., Gates, A. R. & Jones, D. O. Fish food in the deep sea: revisiting the role of large food-falls. PloS one 9, e96016 (2014).

4 Xu, H. et al. Evolution and genetic adaptation of fishes to the deep sea. Cell 188, 1393–1408. e1313 (2025).

5 Zhang, H. et al. The amphipod genome reveals population dynamics and adaptations to hadal environment. Cell 188, 1378–1392. e1318 (2025).

6 Shi, M., Qi, L. & He, L.-S. Comparative Analysis of the Mitochondrial Genome of Galatheanthemum sp. MT-2020 (Actiniaria Galatheanthemidae) From a Depth of 9,462 m at the Mariana Trench. Frontiers in Genetics 13, 854009 (2022).

7 Wolff, T. in Deep Sea Research and Oceanographic Abstracts. 983–1003 (Elsevier).

8 González-Muñoz, R. et al. Sea anemones (Cnidaria: Actiniaria, Corallimorpharia, Ceriantharia, Zoanthidea) from marine shallow-water environments in Venezuela: new records and an updated inventory. Marine Biodiversity Records 9, 1–35 (2016).

9 Fautin, D. G., Malarky, L. & Soberón, J. Latitudinal diversity of sea anemones (Cnidaria: Actiniaria). The Biological Bulletin 224, 89–98 (2013).

10 Reft, A. J. & Daly, M. Morphology, distribution, and evolution of apical structure of nematocysts in hexacorallia. Journal of morphology 273, 121–136 (2012).

11 Campoy, A. N., Rivadeneira, M. M., Hernández, C. E., Meade, A. & Venditti, C. Deep-sea origin and depth colonization associated with phenotypic innovations in scleractinian corals. Nature Communications 14, 7458 (2023).

12 Rodríguez, E. et al. Hidden among sea anemones: the first comprehensive phylogenetic reconstruction of the order Actiniaria (Cnidaria, Anthozoa, Hexacorallia) reveals a novel group of hexacorals. PloS one 9, e96998 (2014).

13 Barroso, F. R. G. et al. Insights into the lifestyle and preservation of Arenactinia ipuensis n. gen. et n. sp.(Anthozoa, Actiniaria) from the Early Silurian (Ipu Formation, Parnaíba Basin, Brazil). Earth History and Biodiversity, 100017 (2025).

14 McMahon, S., Tarhan, L. G. & Briggs, D. E. Decay of the sea anemone Metridium (Actiniaria): implications for the preservation of cnidarian polyps and other soft-bodied diploblast-grade animals. Palaios 32, 388–395 (2017).

15 Seilacher, A., Reif, W.-E. & Westphal, F. Sedimentological, ecological and temporal patterns of fossil Lagerstätten. Philosophical Transactions of the Royal Society of London. B, Biological Sciences 311, 5–24 (1985).

16 Zhang, H., Zhou, Y. & Yang, Z. Genetic adaptations of marine invertebrates to hydrothermal vent habitats. Trends in Genetics (2024).

17 Zhou, Y. et al. Genetic adaptations of sea anemone to hydrothermal environment. Science Advances 9, eadh0474 (2023).

18 Law, S. T. S. et al. The genome of the deep-sea anemone Actinernus sp. contains a mega-array of ANTP-class homeobox genes. Proceedings of the Royal Society B 290, 20231563 (2023).

19 Feng, C. et al. The genome of a new anemone species (Actiniaria: Hormathiidae) provides insights into deep-sea adaptation. Deep Sea Research Part I: Oceanographic Research Papers 170, 103492 (2021).

20 Pimiento, C. et al. The Pliocene marine megafauna extinction and its impact on functional diversity. Nature ecology & evolution 1, 1100–1106 (2017).

21 Melott, A. L. & Bambach, R. K. Analysis of periodicity of extinction using the 2012 geological timescale. Paleobiology 40, 177–196 (2014).

22 Bell-Pedersen, D. et al. Circadian rhythms from multiple oscillators: lessons from diverse organisms. Nature Reviews Genetics 6, 544–556 (2005).

23 Müller, W. E., Schröder, H. C., Pisignano, D., Markl, J. S. & Wang, X. Metazoan circadian rhythm: toward an understanding of a light-based zeitgeber in sponges. Integrative and comparative biology 53, 103–117 (2013).

24 Reppert, S. M. & Weaver, D. R. Coordination of circadian timing in mammals. Nature 418, 935–941 (2002).

25 Panda, S., Hogenesch, J. B. & Kay, S. A. Circadian rhythms from flies to human. Nature 417, 329–335 (2002).

26 Reitzel, A. M., Tarrant, A. M. & Levy, O. Circadian clocks in the cnidaria: environmental entrainment, molecular regulation, and organismal outputs. Integrative and Comparative Biology 53, 118–130 (2013).

27 Simionato, E. et al. Origin and diversification of the basic helix-loop-helix gene family in metazoans: insights from comparative genomics. BMC evolutionary biology 7, 1–18 (2007).

28 Stanton, D., Justin, H. S. & Reitzel, A. M. Step in time: conservation of circadian clock genes in animal evolution. Integrative and Comparative Biology 62, 1503–1518 (2022).

29 Kawamoto, T. et al. A novel autofeedback loop of Dec1 transcription involved in circadian rhythm regulation. Biochemical and biophysical research communications 313, 117–124 (2004).

30. Pinet, P. R. Invitation to oceanography. (Jones & Bartlett Learning, 2006).

31 Izumi, T., Ise, Y., Yanagi, K., Shibata, D. & Ueshima, R. First detailed record of symbiosis between a sea anemone and homoscleromorph sponge, with a description of Tempuractis rinkai gen. et sp. nov.(Cnidaria: Anthozoa: Actiniaria: Edwardsiidae). Zoological Science 35, 188–198 (2018).

32 Beaulieu, S. E. Colonization of habitat islands in the deep sea: recruitment to glass sponge stalks. Deep Sea Research Part I: Oceanographic Research Papers 48, 1121–1137 (2001).

33 Beaulieu, S. Life on glass houses: sponge stalk communities in the deep sea. Marine Biology 138, 803–817 (2001).

34 Young, C. M. in Marine Hard Bottom Communities: Patterns, Dynamics, Diversity, and Change 39–60 (Springer, 2009).

35 Berterame, N. M., Bertagnoli, S., Codazzi, V., Porro, D. & Branduardi, P. Temperature-induced lipocalin (TIL): a shield against stress-inducing environmental shocks in Saccharomyces cerevisiae. FEMS Yeast Research 17, fox056 (2017).

36 Himmel, N. J. & Cox, D. N. Transient receptor potential channels: current perspectives on evolution, structure, function and nomenclature. Proceedings of the Royal Society B 287, 20201309 (2020).

37 Peng, G., Shi, X. & Kadowaki, T. Evolution of TRP channels inferred by their classification in diverse animal species. Molecular phylogenetics and evolution 84, 145–157 (2015).

38 Venkatachalam, K. & Montell, C. TRP channels. Annu. Rev. Biochem. 76, 387–417 (2007).

39 Thiagarajan, V., Byrdin, M., Eker, A. P., Müller, P. & Brettel, K. Kinetics of cyclobutane thymine dimer splitting by DNA photolyase directly monitored in the UV. Proceedings of the National Academy of Sciences 108, 9402–9407 (2011).

40 Sancar, A. Structure and function of DNA photolyase and cryptochrome blue-light photoreceptors. Chemical reviews 103, 2203–2238 (2003).

41 Salih, A., Larkum, A., Cox, G., Kühl, M. & Hoegh-Guldberg, O. Fluorescent pigments in corals are photoprotective. Nature 408, 850–853 (2000).

42 Anthony, K. R. & Fabricius, K. E. Shifting roles of heterotrophy and autotrophy in coral energetics under varying turbidity. Journal of experimental marine biology and ecology 252, 221–253 (2000).

43 Clarke, D. N. et al. Fluorescent proteins generate a genetic color polymorphism and counteract oxidative stress in intertidal sea anemones. Proceedings of the National Academy of Sciences 121, e2317017121 (2024).

44 Strumpen, N. F., Rädecker, N., Pogoreutz, C., Meibom, A. & Voolstra, C. R. High light quantity suppresses locomotion in symbiotic Aiptasia. Symbiosis 86, 293–304 (2022).

45 Pearse, V. B. Modification of sea anemone behavior by symbiotic zooxanthellae: phototaxis. The Biological Bulletin 147, 630–640 (1974).

46 RIEMANN-ZÜRNECK, K. How sessile are sea anemones? A review of free-living forms in the Actiniaria Cnidaria: Anthozoa. Marine Ecology 19, 247–261 (1998).

47 Franklin, E. C., Stat, M., Pochon, X., Putnam, H. M. & Gates, R. D. GeoSymbio: a hybrid, cloud-based web application of global geospatial bioinformatics and ecoinformatics for Symbiodinium–host symbioses. Molecular Ecology Resources 12, 369–373 (2012).

48 Shi, T. et al. Untangling ITS2 genotypes of algal symbionts in zooxanthellate corals. Molecular Ecology Resources 21, 137–152 (2021).

49 Tonk, L., Bongaerts, P., Sampayo, E. M. & Hoegh-Guldberg, O. SymbioGBR: a web-based database of Symbiodinium associated with cnidarian hosts on the Great Barrier Reef. BMC ecology 13, 1–9 (2013).

50 Cui, G. et al. Molecular insights into the Darwin paradox of coral reefs from the sea anemone Aiptasia. Science Advances 9, eadf7108 (2023).

51 Muscatine, L. & Porter, J. W. Reef corals: mutualistic symbioses adapted to nutrient-poor environments. Bioscience 27, 454–460 (1977).

52 Carlisle, J. F., Murphy, G. K. & Roark, A. M. Body size and symbiotic status influence gonad development in Aiptasia pallida anemones. Symbiosis 71, 121–127 (2017).

53 Clayton Jr, W. S. Pedal laceration by the anemone Aiptasia pallida. Marine ecology progress series. Oldendorf 21, 75–80 (1985).

54 Szmant, A. & Gassman, N. The effects of prolonged “bleaching” on the tissue biomass and reproduction of the reef coral Montastrea annularis. Coral reefs 8, 217–224 (1990).

55 Guruacharya, A. Genomic investigation of cnidarian toxin evolution. (2017).

56 Sachkova, M. Y. et al. The birth and death of toxins with distinct functions: a case study in the sea anemone Nematostella. Molecular Biology and Evolution 36, 2001–2012 (2019).

57 Columbus-Shenkar, Y. Y. et al. Dynamics of venom composition across a complex life cycle. Elife 7, e35014 (2018).

58 Jaimes-Becerra, A. et al. “Beyond primary sequence”—proteomic data reveal complex toxins in cnidarian venoms. Integrative and Comparative Biology 59, 777–785 (2019).

59 de Lecea, L. et al. The hypocretins: hypothalamus-specific peptides with neuroexcitatory activity. Proceedings of the National Academy of Sciences 95, 322–327 (1998).

60 Wong, K. K., Ng, S. Y., Lee, L. T., Ng, H. K. & Chow, B. K. Orexins and their receptors from fish to mammals: a comparative approach. General and comparative endocrinology 171, 124–130 (2011).

61 Shick, J. M., Hoffmann, R. J. & Lamb, A. N. Asexual reproduction, population structure, and genotype-environment interactions in sea anemones. American Zoologist 19, 699–713 (1979).

62 Chia, F.-S. in Coelenterate ecology and behavior 261–270 (Springer, 1976).

63 Miramón-Puértolas, P., Pascual-Carreras, E. & Steinmetz, P. R. A population of Vasa2 and Piwi1 expressing cells generates germ cells and neurons in a sea anemone. Nature Communications 15, 8765 (2024).

64 Shao, P.-J., Chiu, Y.-L., Tsai, P.-H. & Shikina, S. Possible germline progenitor cells in extra-gonadal tissues of the sea anemone, Exaiptasia diaphana. Frontiers in Marine Science 10, 1278022 (2023).

65 Ishiguro, K.-i., et al. MEIOSIN directs the switch from mitosis to meiosis in mammalian germ cells. Developmental cell 52, 429–445. e410 (2020).

66 Li, L. et al. The XRN1-regulated RNA helicase activity of YTHDC2 ensures mouse fertility independently of m6A recognition. Molecular Cell 82, 1678–1690. e1612 (2022).

67 Munro, C., Cadis, H., Pagnotta, S., Houliston, E. & Huynh, J.-R. Conserved meiotic mechanisms in the cnidarian Clytia hemisphaerica revealed by Spo11 knockout. Science Advances 9, eadd2873 (2023).

68 Challa, K. et al. Meiosis-specific prophase-like pathway controls cleavage-independent release of cohesin by Wapl phosphorylation. PLoS genetics 15, e1007851 (2019).

69 Lee, M.-H. et al. Multiple functions and dynamic activation of MPK-1 extracellular signal-regulated kinase signaling in Caenorhabditis elegans germline development. Genetics 177, 2039–2062 (2007).

70 Rogacheva, M. V. et al. Mlh1-Mlh3, a meiotic crossover and DNA mismatch repair factor, is a Msh2-Msh3-stimulated endonuclease. Journal of Biological Chemistry 289, 5664–5673 (2014).

71 Santucci-Darmanin, S. et al. The DNA mismatch-repair MLH3 protein interacts with MSH4 in meiotic cells, supporting a role for this MutL homolog in mammalian meiotic recombination. Human molecular genetics 11, 1697–1706 (2002).

72 Gross, M. & Jaenicke, R. Proteins under pressure: the influence of high hydrostatic pressure on structure, function and assembly of proteins and protein complexes. European Journal of Biochemistry 221, 617–630 (1994).

73 Somero, G. N. Adaptations to high hydrostatic pressure. Annual review of physiology 54, 557–577 (1992).

74 Wang, K. et al. Morphology and genome of a snailfish from the Mariana Trench provide insights into deep-sea adaptation. Nature ecology & evolution 3, 823–833 (2019).

75 Yancey, P. H. Adaptations to hydrostatic pressure in protein structure and organic osmolytes in deep-sea animals. High Pressure Bioscience and Biotechnology 1, 90–95 (2007).

76 Siebenaller, J. & Somero, G. N. Pressure-adaptive differences in lactate dehydrogenases of congeneric fishes living at different depths. Science 201, 255–257 (1978).

77 Chen, M., Markham, J. E. & Cahoon, E. B. Sphingolipid Δ8 unsaturation is important for glucosylceramide biosynthesis and low-temperature performance in Arabidopsis. The plant journal 69, 769–781 (2012).

78 Mashima, R., Okuyama, T. & Ohira, M. Biosynthesis of long chain base in sphingolipids in animals, plants and fungi. Future science OA 6, FSO434 (2020).

79 Endo, K., Akiyama, T., Kobayashi, S. & Okada, M. Degenerative spermatocyte, a novel gene encoding a transmembrane protein required for the initiation of meiosis in Drosophila spermatogenesis. Molecular and General Genetics MGG 253, 157–165 (1996).

80 Ternes, P., Franke, S., Zähringer, U., Sperling, P. & Heinz, E. Identification and characterization of a sphingolipid Δ4-desaturase family. Journal of Biological Chemistry 277, 25512–25518 (2002).

81 Xu, W. et al. Chromosome-level genome assembly of hadal snailfish reveals mechanisms of deep-sea adaptation in vertebrates. Elife 12, RP87198 (2023).

82 Murrell, D. J. A global envelope test to detect non-random bursts of trait evolution. Methods in Ecology and Evolution 9, 1739–1748 (2018).

83 Bollback, J. P. SIMMAP: stochastic character mapping of discrete traits on phylogenies. BMC bioinformatics 7, 1–7 (2006).

84 Chen, S., Zhou, Y., Chen, Y. & Gu, J. fastp: an ultra-fast all-in-one FASTQ preprocessor. Bioinformatics 34, i884–i890 (2018).

85 Luo, R. et al. SOAPdenovo2: an empirically improved memory-efficient short-read de novo assembler. Gigascience 1, 2047–2217X-2041-2018 (2012).

86 Ruan, J. & Li, H. Fast and accurate long-read assembly with wtdbg2. Nature methods 17, 155–158 (2020).

87 Hu, J., Fan, J., Sun, Z. & Liu, S. NextPolish: a fast and efficient genome polishing tool for long-read assembly. Bioinformatics 36, 2253–2255 (2020).

88 Cheng, H. et al. Haplotype-resolved assembly of diploid genomes without parental data. Nature biotechnology 40, 1332–1335 (2022).

89 Guan, D. et al. Identifying and removing haplotypic duplication in primary genome assemblies. Bioinformatics 36, 2896–2898 (2020).

90 Kajitani, R. et al. Efficient de novo assembly of highly heterozygous genomes from whole-genome shotgun short reads. Genome research 24, 1384–1395 (2014).

91 Alonge, M. et al. Automated assembly scaffolding elevates a new tomato system for high-throughput genome editing. BioRxiv, 2021.2011. 2018.469135 (2021).

92 Simão, F. A., Waterhouse, R. M., Ioannidis, P., Kriventseva, E. V. & Zdobnov, E. M. BUSCO: assessing genome assembly and annotation completeness with single-copy orthologs. Bioinformatics 31, 3210–3212 (2015).

93 Allio, R. et al. MitoFinder: Efficient automated large-scale extraction of mitogenomic data in target enrichment phylogenomics. Molecular ecology resources 20, 892–905 (2020).

94 Bernt, M. et al. MITOS: improved de novo metazoan mitochondrial genome annotation. Molecular phylogenetics and evolution 69, 313–319 (2013).

95 Bushmanova, E., Antipov, D., Lapidus, A. & Prjibelski, A. D. rnaSPAdes: a de novo transcriptome assembler and its application to RNA-Seq data. GigaScience 8, giz100 (2019).

96 Saha, S., Bridges, S., Magbanua, Z. V. & Peterson, D. G. Empirical comparison of ab initio repeat finding programs. Nucleic acids research 36, 2284–2294 (2008).

97 Chen, N. Using Repeat Masker to identify repetitive elements in genomic sequences. Current protocols in bioinformatics 5, 4.10. 11–14.10. 14 (2004).

98 Tempel, S. Using and understanding RepeatMasker. Mobile genetic elements: protocols and genomic applications, 29–51 (2012).

99 Bao, W., Kojima, K. K. & Kohany, O. Repbase Update, a database of repetitive elements in eukaryotic genomes. Mobile Dna 6, 1–6 (2015).

100 Benson, G. Tandem repeats finder: a program to analyze DNA sequences. Nucleic acids research 27, 573–580 (1999).

101 Li, M. et al. Genomes of Two Flying Squid Species Provide Novel Insights into Adaptations of Cephalopods to Pelagic Life. Genomics, Proteomics & Bioinformatics 20, 1053–1065 (2022).

102 Hu, M. et al. A chromosome-level genome of the striated frogfish (Antennarius striatus). Scientific Data 11, 654 (2024).

103 Stanke, M., Diekhans, M., Baertsch, R. & Haussler, D. Using native and syntenically mapped cDNA alignments to improve de novo gene finding. Bioinformatics 24, 637–644 (2008).

104 Kent, W. J. BLAT—the BLAST-like alignment tool. Genome research 12, 656–664 (2002).

105 Birney, E., Clamp, M. & Durbin, R. GeneWise and genomewise. Genome research 14, 988–995 (2004).

106 Haas, B. J., Zeng, Q., Pearson, M. D., Cuomo, C. A. & Wortman, J. R. Approaches to fungal genome annotation. Mycology 2, 118–141 (2011).

107 Haas, B. J. et al. Automated eukaryotic gene structure annotation using EVidenceModeler and the Program to Assemble Spliced Alignments. Genome biology 9, 1–22 (2008).

108 Jones, P. et al. InterProScan 5: genome-scale protein function classification. Bioinformatics 30, 1236–1240 (2014).

109 Katoh, K., Misawa, K., Kuma, K. i. & Miyata, T. MAFFT: a novel method for rapid multiple sequence alignment based on fast Fourier transform. Nucleic acids research 30, 3059–3066 (2002).

110 Capella-Gutiérrez, S., Silla-Martínez, J. M. & Gabaldón, T. trimAl: a tool for automated alignment trimming in large-scale phylogenetic analyses. Bioinformatics 25, 1972–1973 (2009).

111 Minh, B. Q. et al. IQ-TREE 2: new models and efficient methods for phylogenetic inference in the genomic era. Molecular biology and evolution 37, 1530–1534 (2020).

112 Zhang, C., Rabiee, M., Sayyari, E. & Mirarab, S. ASTRAL-III: polynomial time species tree reconstruction from partially resolved gene trees. BMC bioinformatics 19, 15–30 (2018).

113 Yang, Z. PAML 4: phylogenetic analysis by maximum likelihood. Molecular biology and evolution 24, 1586–1591 (2007).

114 Pease, J. B., Brown, J. W., Walker, J. F., Hinchliff, C. E. & Smith, S. A. Quartet Sampling distinguishes lack of support from conflicting support in the green plant tree of life. American journal of botany 105, 385–403 (2018).

115 Sayyari, E., Whitfield, J. B. & Mirarab, S. DiscoVista: interpretable visualizations of gene tree discordance. Molecular Phylogenetics and Evolution 122, 110–115 (2018).

116 Minh, B. Q., Hahn, M. W. & Lanfear, R. New methods to calculate concordance factors for phylogenomic datasets. Molecular biology and evolution 37, 2727–2733 (2020).

117 Galtier, N. & Daubin, V. Dealing with incongruence in phylogenomic analyses. Philosophical Transactions of the Royal Society B: Biological Sciences 363, 4023–4029 (2008).

118 Edwards, S. V. Is a new and general theory of molecular systematics emerging? Evolution 63, 1–19 (2009).

119 Xu, B. & Yang, Z. Challenges in species tree estimation under the multispecies coalescent model. Genetics 204, 1353–1368 (2016).

120 Wen, D. & Nakhleh, L. Coestimating reticulate phylogenies and gene trees from multilocus sequence data. Systematic biology 67, 439–457 (2018).

121 Rhodes, J. A., Baños, H., Mitchell, J. D. & Allman, E. S. MSCquartets 1.0: quartet methods for species trees and networks under the multispecies coalescent model in R. Bioinformatics 37, 1766–1768 (2021).

122 Li, H. & Durbin, R. Inference of human population history from individual whole-genome sequences. Nature 475, 493–496 (2011).

123 Li, H. et al. The sequence alignment/map format and SAMtools. bioinformatics 25, 2078–2079 (2009).

124 Sanderson, M. J. r8s: inferring absolute rates of molecular evolution and divergence times in the absence of a molecular clock. Bioinformatics 19, 301–302 (2003).

125 Emms, D. M. & Kelly, S. OrthoFinder: solving fundamental biases in whole genome comparisons dramatically improves orthogroup inference accuracy. Genome biology 16, 1–14 (2015).

126 Stamatakis, A. RAxML version 8: a tool for phylogenetic analysis and post-analysis of large phylogenies. Bioinformatics 30, 1312–1313 (2014).

127 Buchfink, B., Xie, C. & Huson, D. H. Fast and sensitive protein alignment using DIAMOND. Nature methods 12, 59–60 (2015).

128. Li, H. Aligning sequence reads, clone sequences and assembly contigs with BWA-MEM. arXiv preprint arXiv:1303. 3997 (2013).

129 Li, Y. et al. Origin and stepwise evolution of vertebrate lungs. Nature Ecology & Evolution, 1–20 (2025).

130 Ranwez, V., Douzery, E. J., Cambon, C., Chantret, N. & Delsuc, F. MACSE v2: toolkit for the alignment of coding sequences accounting for frameshifts and stop codons. Molecular biology and evolution 35, 2582–2584 (2018).

131 Bu, D. et al. KOBAS-i: intelligent prioritization and exploratory visualization of biological functions for gene enrichment analysis. Nucleic acids research 49, W317–W325 (2021).

132 Kim, D., Paggi, J. M., Park, C., Bennett, C. & Salzberg, S. L. Graph-based genome alignment and genotyping with HISAT2 and HISAT-genotype. Nature biotechnology 37, 907–915 (2019).

133 Pertea, M., Kim, D., Pertea, G. M., Leek, J. T. & Salzberg, S. L. Transcript-level expression analysis of RNA-seq experiments with HISAT, StringTie and Ballgown. Nature protocols 11, 1650–1667 (2016).

134 Gornik, S. G. et al. Photoreceptor diversification accompanies the evolution of Anthozoa. Molecular biology and evolution 38, 1744–1760 (2021).

135 Bernardet, C., Tambutté, E., Techer, N., Tambutté, S. & Venn, A. Ion transporter gene expression is linked to the thermal sensitivity of calcification in the reef coral Stylophora pistillata. Scientific Reports 9, 18676 (2019).

136 Dhar, A. K., Bowers, R. M., Licon, K. S., Veazey, G. & Read, B. Validation of reference genes for quantitative measurement of immune gene expression in shrimp. Molecular Immunology 46, 1688–1695 (2009).

137 Zhang, S. et al. Identification and validation of reference genes for normalization of gene expression analysis using qRT-PCR in Helicoverpa armigera (Lepidoptera: Noctuidae). Gene 555, 393–402 (2015).

138 Kosakyan, A. et al. Selection of suitable reference genes for gene expression studies in myxosporean (Myxozoa, Cnidaria) parasites. Scientific reports 9, 15073 (2019).

139 Lehnert, E. M. et al. Extensive differences in gene expression between symbiotic and aposymbiotic cnidarians. G3: Genes, Genomes, Genetics 4, 277-295 (2014).

140 Silvestro, D. et al. Early arrival and climatically-linked geographic expansion of New World monkeys from tiny African ancestors. Systematic Biology 68, 78–92 (2019).

141 Farina, B. M., Faurby, S. & Silvestro, D. Dollo meets Bergmann: morphological evolution in secondary aquatic mammals. Proceedings of the Royal Society B 290, 20231099 (2023).

142 Rolland, J. et al. The impact of endothermy on the climatic niche evolution and the distribution of vertebrate diversity. Nature ecology & evolution 2, 459–464 (2018).

143 Harmon, L. J., Weir, J. T., Brock, C. D., Glor, R. E. & Challenger, W. GEIGER: investigating evolutionary radiations. Bioinformatics 24, 129–131 (2008).

144 Hendry, A. P. & Kinnison, M. T. Perspective: the pace of modern life: measuring rates of contemporary microevolution. Evolution 53, 1637–1653 (1999).

145 Darwin, C. On the origin of species by means of natural selection, or, the preservation of favoured races in the struggle for life. Harvard Botany Libraries--Biodiversity Heritage Library digitization project (1859).

146 Haldane, J. B. S. in Selected Genetic Papers of JBS Haldane (Routledge Revivals) 127–132 (Routledge, 2022).

147 Wang, W. et al. The China national GeneBank sequence archive (CNSA) 2024 update. Horticulture Research, uhaf036 (2025).

